# Terminal Modification, Sequence, and Length Determine Small RNA Stability in Animals

**DOI:** 10.1101/2020.09.08.287979

**Authors:** Ildar Gainetdinov, Cansu Colpan, Katharine Cecchini, Paul Albosta, Karina Jouravleva, Joel Vega-Badillo, Yongjin Lee, Deniz M. Özata, Phillip D. Zamore

## Abstract

In animals, piRNAs, siRNAs, and miRNAs silence transposons, fight viral infections, and regulate gene expression. piRNA biogenesis concludes with 3′ terminal trimming and 2′-*O*-methylation. Both trimming and methylation influence piRNA stability. Here, we report that trimming and methylation protect mouse piRNAs from different decay mechanisms. In the absence of 2′-*O*-methylation, mouse piRNAs with extensive complementarity to long RNAs become unstable. In flies, 2′-*O*-methylation similarly protects both piRNAs and siRNAs from complementarity-dependent destabilization. Animal miRNAs are unmethylated, and complementarity-dependent destabilization helps explain differences in miRNA decay rates in both mice and flies. In contrast, trimming protects mouse piRNAs from a separate degradation pathway unaffected by target complementarity but sensitive to the 3′ terminal, untrimmed sequence. Because distinct sets of mouse piRNAs are protected by each of these mechanisms, loss of both trimming and 2′-*O*-methylation causes the piRNA pathway to collapse, demonstrating that these two small RNA modifications collaborate to stabilize piRNAs.

**Highlights:**

- 2′-*O*-methylation protects mouse and fly piRNAs from complementarity-dependent decay
- 2′-*O*-methylation protects fly siRNAs with extensive complementarity to long RNAs
- Complementarity to long RNAs predicts the half-life of fly and mouse miRNAs
- Mouse pre-piRNA decay reflects both pre-piRNA sequence and PIWI protein identity

## INTRODUCTION

In animals, three classes of small silencing RNAs direct Argonaute proteins to target RNAs. MicroRNAs (miRNAs) regulate host mRNAs (Bartel, 2018). Small interfering RNAs (siRNAs) target host, transposon and viral transcripts (Carthew and Sontheimer, 2009). PIWI-interacting RNAs (piRNAs) defend the genome against transposable elements and, in some animals, also regulate gene expression or fight viral infection (Huang et al., 2017; Czech et al., 2018; Yamashiro and Siomi, 2018; Ozata et al., 2019). Although their sequences, lengths and genomic origins vary, piRNAs guide members of the PIWI clade of Argonaute proteins (PIWI proteins) in nearly all animals, including sponges, cnidarians, arthropods, nematodes and chordates (Aravin et al., 2006; Girard et al., 2006; Lau et al., 2006; Vagin et al., 2006; Grivna et al., 2006; Saito et al., 2006; Houwing et al., 2007; Grimson et al., 2008; Das et al., 2008; Batista et al., 2008; Juliano et al., 2014; Lim et al., 2014; Lewis et al., 2018).

Unlike miRNA and siRNA biogenesis, piRNA production begins with long single-stranded, not double-stranded, RNA (Vagin et al., 2006). piRNA precursors are transcribed by RNA polymerase II from dedicated genomic loci called piRNA clusters (Brennecke et al., 2007; Aravin et al., 2006; Girard et al., 2006; Li et al., 2013; Fu et al., 2018; Özata et al., 2020). PIWI proteins guided by pre-existing piRNAs cleave these precursors, creating 5′ monophosphorylated pre-pre-piRNAs on which piRNA biogenesis initiates (Wang et al., 2014; Han et al., 2015; Gainetdinov et al., 2018). The 5′ monophosphate is required to load the pre-pre-piRNA into a PIWI protein, a process believed to occur in a structure called nuage (Kawaoka et al., 2011; Cora et al., 2014; Wang et al., 2014; Mohn et al., 2015; Han et al., 2015; Matsumoto et al., 2016; Yamaguchi et al., 2020). After relocalization of the PIWI-bound pre-pre-piRNA to the outer mitochondrial membrane, the endonuclease PLD6 (Zucchini in *Drosophila melanogaster*) cleaves the pre-pre-piRNAs 3′ to the footprint of the PIWI protein, releasing a PIWI-bound pre-piRNA and a new 5′ monophosphorylated pre-pre-piRNA that can bind yet another PIWI protein (Haase et al., 2010; Ipsaro et al., 2012; Nishimasu et al., 2012; Han et al., 2015; Mohn et al., 2015; Homolka et al., 2015; Gainetdinov et al., 2018; Ge et al., 2019; Munafò et al., 2019; Ishizu et al., 2019; Izumi et al., 2020). Successive cycles of pre-pre-piRNA binding by PIWI proteins and cleavage by PLD6 convert the original piRNA precursor transcript into phased, tail-to-head strings of pre-piRNAs.

In the penultimate step in piRNA biogenesis, the 3′-to-5′ exoribonuclease PNLDC1 in *Mus musculus*, Trimmer in most arthropods, or PARN1 in *Caenorhabditis elegans* establishes the mature length of the piRNA, which reflects the footprint of the specific PIWI protein to which the piRNA is bound (Tang et al., 2016; Izumi et al., 2016; Zhang et al., 2017; Ding et al., 2017; Nishimura et al., 2018; Gainetdinov et al., 2018). (In flies and likely other members of the Brachycera suborder of Diptera, the miRNA-trimming endoribonuclease, Nibbler, takes the place of PNLDC1 to trim a subset of piRNAs; Feltzin et al., 2015; Wang et al., 2016; Hayashi et al., 2016; Han et al., 2011; Liu et al., 2011). piRNA biogenesis concludes when the *S*-adenosylmethionine-dependent methyltransferase HENMT1 in mice, Hen1 in arthropods, and HENN1 in worms modifies the 2′ hydroxyl at the 3′ end of the piRNA (Saito et al., 2007; Horwich et al., 2007; Kirino and Mourelatos, 2007; Kamminga et al., 2010; Montgomery et al., 2012; Billi et al., 2012; Kamminga et al., 2012; Lim et al., 2015; Svendsen et al., 2019). *Pnldc1*^−/−^ mutant mice accumulate untrimmed pre-piRNAs bound to PIWI proteins (Ding et al., 2017; Zhang et al., 2017; Nishimura et al., 2018; Gainetdinov et al., 2018). The abundance of these piRNA intermediates is 50–70% lower than in wild-type (Gainetdinov et al., 2018). Conversely, *Henmt1*^−/−^ mutant males trim their piRNAs but cannot methylate their 3′ termini. Failure to methylate piRNAs halves the abundance of unmethylated piRNAs (Lim et al., 2015). The molecular defects in these mutants suggest that both trimming and 2′-*O*-methylation play a role in stabilizing piRNAs.

Dicer enzymes set the mature length of animal miRNAs and siRNAs, and they are typically stable without further modification (Bartel, 2018). However, most arthropod siRNAs are 2′-*O*-methylated (Pélisson et al., 2007; Lewis et al., 2018; Fu et al., 2018); in flies, siRNAs are unstable in the absence of 2′-*O*-methylation (Ameres et al., 2010).

Here, we report that 3′ terminal 2′-*O*-methylation and 3′-to-5′ trimming protect mouse piRNAs against distinct degradation mechanisms. In the absence of 2′-*O*-methylation, piRNAs with extensive complementarity to long RNAs are destroyed. We provide evidence that 2′-*O*-methylation similarly blocks complementarity-dependent destabilization for both piRNAs and siRNAs in flies. Complementarity-dependent destabilization also helps explain differences in decay rates among miRNAs in both mouse and fly cell lines. In contrast, long complementary RNAs do not destabilize untrimmed mouse pre-piRNAs. Instead, both PIWI protein identity and the presence of oligouridine or oligoguanine tracts in the untrimmed sequence of a pre-piRNA correlate with instability. In *Pnldc1*^*em1/em1*^; *Henmt1*^*em1/em1*^ double-mutant males, which can neither trim nor methylate piRNAs, the piRNA pathway collapses. Our data demonstrate that piRNA trimming and methylation collaborate to stabilize piRNAs: the double-mutant mice make sixfold fewer piRNAs, and spermatogenesis arrests at the pachytene stage of meiosis. The reduction in piRNA abundance derepresses both mRNA and transposon transcript targets. We propose that by decreasing the degradation rate of piRNAs, methylation and trimming maintain the high steady-state abundance that piRNAs require to repress their RNA targets.

## RESULTS

### 2′-*O*-methylation Inhibits Complementarity-Dependent Destabilization of Mouse piRNAs

miRNAs with extensively complementary targets are unstable—a phenomenon termed Target RNA-Directed miRNA Degradation (TDMD; Cazalla et al., 2010; Ameres et al., 2010; Baccarini et al., 2011; Libri et al., 2012; Marcinowski et al., 2012; Rüegger and Großhans, 2012; Lee et al., 2013; de la Mata et al., 2015; Bitetti et al., 2018; Kleaveland et al., 2018; Ghini et al., 2018; Sheu-Gruttadauria et al., 2019a; Zhang et al., 2019). In most animals, miRNAs bear a 2′-hydroxyl at their 3′ terminus. In *D. melanogaster*, the subset of miRNAs loaded into the siRNA-guided protein Ago2 are 2′-*O*-methylated, and this modification protects Ago2-bound miRNAs against TDMD (Ameres et al., 2010). miRNAs in the sea anemone *Nematostella vectensis* often target transcripts through near-perfect complementarity (Moran et al., 2014). All *N. vectensis* miRNAs are 2′-*O*-methylated to some extent, and depletion of the *N. vectensis* homolog of the methyltransferase HENMT1 reduces miRNA stability (Moran et al., 2014; Modepalli et al., 2018).

Does 2′-*O*-methylation also protect mouse piRNAs from a degradation mechanism dependent on extensive complementarity to long RNAs? We generated a mouse mutant lacking functional HENMT1 protein, *Henmt1*^*em1Pdz/em1Pdz*^ (henceforth, *Henmt1^em1/em1^*). piRNAs in *Henmt1*^*em1/em1*^ mice lack 2′-*O*-methylation at their 3′ termini, and the abundance of most piRNAs decreases, albeit to varying extents, ranging from 0 to 100% of C57BL/6 levels (Figure S1A). If complementarity-dependent destabilization explains piRNA degradation in the absence of 2′-*O*-methylation, unstable piRNAs are expected to have more abundant and higher affinity complementary sites in the transcriptome than stable piRNAs.

We sequenced long transcripts from FACS-sorted mouse germ cells, and, for each piRNA, calculated the cumulative concentration of all sites with 5 to 11 contiguously complementary nucleotides present in the transcriptome, [complementary sites]_*total*_ (Figure 1A). RNA duplexes shorter than 5 nt are not expected to be stable (Duchesne, 1973); 11 nucleotides was the longest stretch for which the majority of piRNAs contained at least one complementary site in the transcriptome. We iteratively determined the [complementary sites]_*total*_ for stretches of complementarity starting at each piRNA nucleotide from g2 to g25 (Figure 1A). Using the equilibrium assumption allows calculation of the fraction of each piRNA region bound to its complementary sites (see STAR Methods):

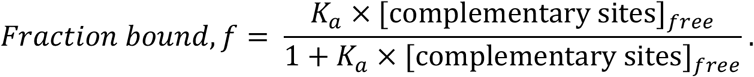

**Figure 1.**
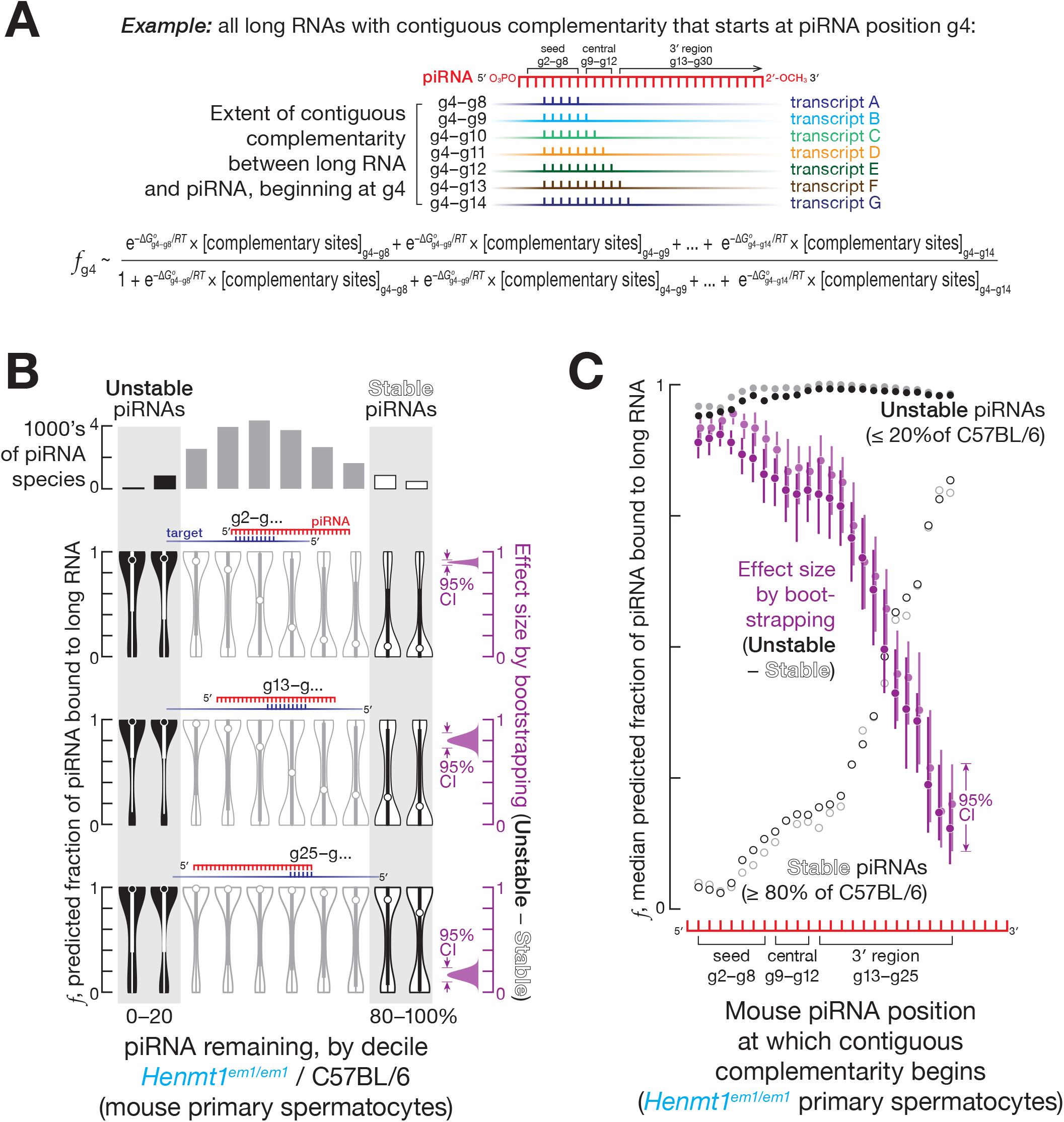
Mouse piRNA 2′-*O*-methylation Protects from Destabilization Dependent on Complementarity to Long RNAs. (A) Strategy to estimate the fraction of piRNA bound to complementary sites in long RNAs. (B) Predicted fraction for different regions of mouse pachytene piRNAs bound to complementary sites in the transcriptome for a representative experiment from FACS-purified *Henmt1*^*em1/em1*^ primary spermatocytes. The 95% confidence interval for the effect size of median difference was calculated with 10,000 bootstrapping iterations. (C) Analysis of mouse pachytene piRNAs from FACS-purified primary spermatocytes showing the median predicted fraction bound for complementarity starting at piRNA positions g2–g25 for stable piRNAs (≥ 80% of C57BL/6 levels in *Henmt1*^*em1/em1*^) for unstable piRNAs (≤ 20% of C57BL/6 levels in *Henmt1*^*em1/em1*^), as well as the difference between the two (i.e., unstable piRNAs − stable piRNAs) for two independent experiments (shown in different shades of the same color). The 95% confidence interval for the effect size of median difference was calculated with 10,000 bootstrapping iterations.

We used two approximations to compare fraction bound (*f*) among piRNAs. First, we presumed that the rank order of [complementary sites]_*free*_ for different piRNAs can be approximated by the rank order of [complementary sites]_*total*_ for those piRNAs. Second, because the binding affinity of different regions of a piRNA bound to a PIWI protein remains unknown, we used the predicted Gibbs free energy (Δ*G*°) of base pairing between two RNA strands at 33°C (Kandeel and Swerdloff, 1988) to estimate the rank order of affinities of different piRNA regions for complementary sites 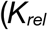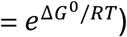. The fraction of an individual piRNA species bound to complementary sites can therefore be approximated as:

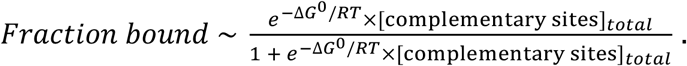

For each nucleotide of each piRNA, we estimated the fraction bound accounting for the contribution of 5–11 nt stretches of complementarity starting at the same nucleotide: e.g., the fraction bound for g4 includes sites complementary to piRNA nucleotides g4–g8, g4–g9, g4–g10, g4–g11, g4–g12, g4–g13, and g4–g14 (Figure 1A).

For complementarity sites starting at g2, g9, and g13, the median fraction bound was ≥ 0.9 for *unstable* piRNAs, those whose steady-state abundance in *Henmt1*^*em1/em1*^ was reduced to ≤ 20% of C57BL/6 levels (Figures 1B and 1C). In contrast, the median fraction bound was ≤ 0.3 for *stable* piRNAs, those whose steady-state abundance remained ≥ 80% of C57BL/6 levels (Figures 1B and 1C). Thus, the predicted fraction bound was higher for unstable piRNAs compared to stable piRNAs for sites complementary to piRNA seed, central, and 3′ regions (Figures 1B, 1C, S1B and S1C): e.g., in primary spermatocytes, for complementary sites starting at nucleotide g2, the difference in median fraction bound between unstable and stable piRNAs was ~0.9 (95% confidence interval [CI]: 0.87–0.92; all CIs calculated by bootstrapping; Figure 1B). These analyses suggest that piRNAs with extensively complementary sites in the transcriptome are more likely to be degraded in the absence of 2′-*O*-methylation.

Pairing to the seed region (g2–g7) of a miRNA is required to trigger TDMD (Cazalla et al., 2010; Ameres et al., 2010; Sheu-Gruttadauria et al., 2019a). In contrast, seed complementarity was not needed to trigger complementarity-dependent destabilization of mouse piRNAs: complementary sites starting at piRNA position g13 were as effective in promoting complementarity-dependent destabilization as sites bearing complementarity to both the seed (g2–g7) and a region of extensive complementarity beginning at g13. For both site types, the estimated fraction of a piRNA bound to its complementary sites was 10–100 times greater for unstable piRNAs compared to stable piRNAs (Figures 1B and S1D). That is, complementary sites starting at position g13 promoted piRNA loss, and targets combining such sites with seed complementarity caused no additional destabilization. Together, these data suggest that complementarity-dependent piRNA destabilization in the absence of 2′-*O*-methylation does not require pairing to the piRNA seed sequence.

Unmethylated piRNAs were most unstable when the 3′ terminal nucleotides of a piRNA were not paired to the complementary long RNA. We examined contiguous matches of different lengths and, for each length, identified the base pairing pattern associated with the largest difference in the fraction bound between stable and unstable piRNAs (Figure S2A). For all contiguous stretches of complementarity, the base pairing pattern that best explained piRNA instability did not extend beyond position g24 (Figure S2A). Because most mouse piRNAs are ≥ 26 nt long, we conclude that an unmethylated piRNA is most unstable when its 3′ terminal nucleotides are unpaired. Perhaps unpaired terminal nucleotides allow endo- or 3′-to-5′ exo-ribonucleases to access the piRNA.

Sufficiently efficient, piRNA-directed, PIWI-catalyzed target cleavage might protect an unmethylated piRNA from complementarity-dependent destabilization. Our data suggest that target cleavage is not protective. Extensive pairing between a long RNA and a piRNA starting at nucleotide g2 destabilizes the piRNA in the absence of 2′-*O*-methylation (Figures 1B, 1C, S1B, and S1C). Such a pattern of pairing is expected to permit target cleavage (Reuter et al., 2011; Zhang et al., 2015; Goh et al., 2015; Wu et al., 2020). To identify piRNAs that direct target RNA cleavage, we sequenced long 5′ monophosphorylated RNA from C57BL/6 primary spermatocytes, and identified candidate piRNA-directed 3′ cleavage products: long 5′ monophosphorylated RNAs predicted to be produced by PIWI-catalyzed cleavage directed by pairing to piRNA nucleotides g2–g14 (Reuter et al., 2011; Wang et al., 2014; Zhang et al., 2015; Goh et al., 2015), i.e., 5′ monophosphorylated RNAs whose first nine nucleotides plus four nucleotides immediately 5′ to the cleavage site are fully complementary to piRNA nucleotides g2–g14 (Figure S2B). piRNAs with and without detectable 3′ cleavage products were similarly unstable: the median unmethylated piRNA abundance was ~45% of C57BL/6 levels for piRNAs with 3′ cleavage products compared to ~48% of C57BL/6 levels for piRNAs with no detectable 3′ cleavage products (Figure S2B). Thus, target cleavage has little impact on complementarity-dependent piRNA destabilization, likely because the concentration of sites whose extent of complementarity is sufficient to elicit complementarity-dependent destabilization but not to direct target cleavage is much greater than the concentration of sites that can both induce destabilization and be cleaved. We conclude that, in the mouse testis, 3′ terminal 2′-*O*-methylation protects piRNAs from degradation elicited by complementary long RNAs.

### 2′-*O*-methylation Inhibits Complementarity-Dependent Destabilization of Fly piRNAs

Does 2′-*O*-methylation protect piRNAs from complementarity-dependent destabilization in other animals? We sequenced small and long RNAs from the ovaries of control (*w*^*1118*^) and *hen1*^*f00810*^ mutant *D. melanogaster* (Horwich et al., 2007): the majority of unmethylated piRNAs in *hen1*^*f00810*^ fly ovaries were ≤ 20% of control (Figures S3A). Using the equilibrium approach we developed for mouse piRNAs (Figure 1A), we estimated the fraction of each fly piRNA bound to contiguously complementary sites in the transcriptome at 25°C. As in mice, the fraction of unstable piRNAs bound to complementary long RNAs was greater than that of stable piRNAs: e.g., for complementary sites starting at piRNA nucleotide g14, the difference in the median of the predicted fraction bound between unstable and stable piRNAs was ~0.74 (95% CI: 0.11–0.82; Figure 2A). In contrast to mouse piRNAs for which complementarity-dependent destabilization was triggered by long RNAs complementary to any region of piRNA (Figures 1B, 1C, S1B and S1C), fly piRNAs were destabilized by transcripts with extensive complementarity to either the central or 3′ regions of the piRNA: i.e., complementary sites beginning at nucleotides g9–g16 (Figures 2A and S3B).

**Figure 2.**
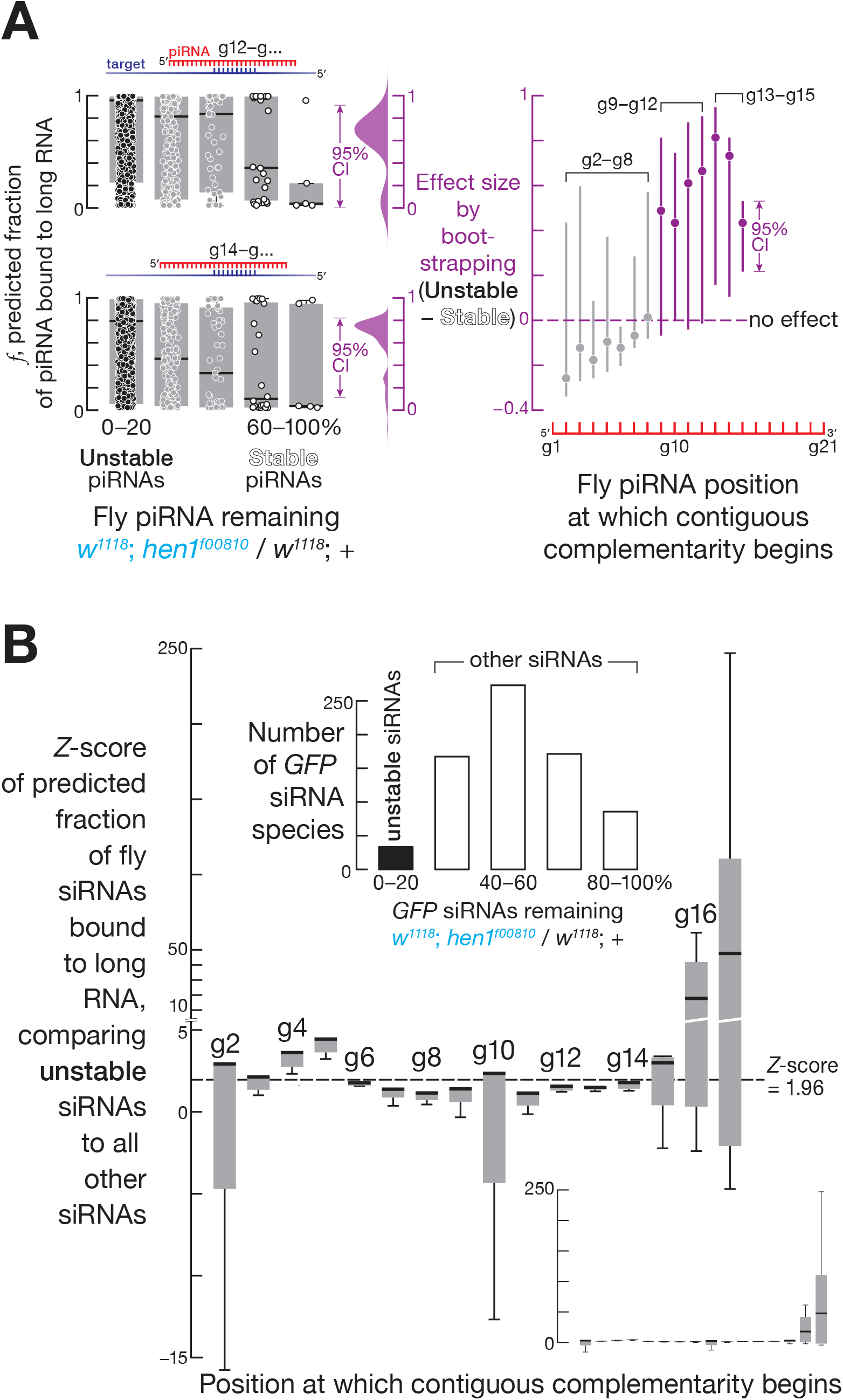
Fly piRNA and siRNA 2′-*O*-methylation Protects against Destabilization Promoted by Complementarity to Long RNAs. (A) Left, mean (*n* = 2) predicted fraction for different regions of fly piRNAs bound to complementary sites in the transcriptome. Right, the difference (i.e., unstable piRNAs - stable piRNAs) between the median predicted fraction bound for complementarity starting at piRNA positions g2–g15 for stable (≥ 80% of *w*^*1118*^ levels in *hen1*^*f00810*^) and unstable piRNAs (≤ 20% of *w*^*1118*^ levels in *hen1*^*f00810*^). Grey: effect size for patterns of complementarity for which the median predicted fraction bound of adjacent quintiles failed to decrease monotonically in the unstable-to-stable direction in Figure S3B. The 95% confidence interval for the effect size of median difference was calculated with 10,000 bootstrapping iterations. (B) Change in abundance (top) of GFP-derived siRNAs in heads from *hen1*^*f00810*^ flies, and *Z*-scores of predicted fraction of different regions of unstable siRNAs bound to complementary sites in the transcriptome.

Like unmethylated piRNAs in mice, unmethylated fly piRNAs did not require pairing with the seed sequence to become unstable when bound to a complementary long RNA: e.g., for complementary sites starting at piRNA nucleotide g14 the estimated fraction bound was ~10 times higher for unstable piRNAs compared to stable piRNAs both when only 3′ region was required to pair (Figure S3B) and when both seed and 3′ region pairing were required (Figure S3C).

We conclude that, as in mice, 3′ terminal 2′-*O*-methylation in flies protects piRNAs from complementarity-dependent destabilization. Unlike mice, whose unmethylated piRNAs are destabilized by long RNAs with a sufficiently long stretch of complementarity to any part of the piRNA, fly piRNAs are destabilized only by complementarity to the central or 3′ regions.

### 2′-*O*-methylation Protects Fly siRNAs from Complementarity-Dependent Destabilization

The 3′ termini of siRNAs are 2′-*O*-methylated in most insect orders, including Hymenoptera, Coleoptera, and Diptera, but not Lepidoptera (Pélisson et al., 2007; Lewis et al., 2018; Fu et al., 2018). In flies, siRNAs derive from long hairpin RNAs encoded in the genome, double-stranded RNAs from viral replication intermediates, transposon transcripts, or convergent transcription (Wang et al., 2006; Galiana-Arnoux et al., 2006; Czech et al., 2008; Ghildiyal et al., 2008; Kawamura et al., 2008; Okamura et al., 2008b; Okamura et al., 2008a; Lau et al., 2009). Fly siRNAs therefore target transposon, viral, and endogenous transcripts via extensive or complete complementarity and are expected to be subject to complementarity-dependent destabilization when unmethylated. Indeed, endo-siRNA abundance declines in *hen1*^*f00810*^ mutant flies (Ameres et al., 2010).

To determine if complementarity-dependent destabilization can explain the instability of unmethylated siRNAs, we used an eye-specific Gal4 driver, *P*(longGMR-GAL4)*3*, to promote transcription of the transgene *P*(UAS-GFP.dsRNA.R)*142*, which produces a 1,440-nt inverted-repeat RNA corresponding to the entire *GFP* coding sequence and measured the abundance of *GFP* siRNAs in control (*w*^*1118*^) and *hen1*^*f00810*^ mutant flies. The abundance of siRNAs derived from the *GFP* inverted repeat transcript was halved in *hen1*^*f00810*^ heads (Figure S3D). As we observed for piRNAs in mice and flies, the abundance of some GFP-siRNAs was unaltered by loss of 3′ terminal 2′-*O*-methylation, while other GFP-siRNAs became unstable (Figure 2B).

We used the equilibrium approach (Figure 1A) to estimate the fraction of each siRNA bound to various contiguously complementary sites in long RNAs. The majority of the predicted fraction bound data clustered near ~1, primarily due to the high predicted binding energies of GFP-derived siRNAs at 25°C (Figure S3E; GC content of GFP sequence is ~62% compared to ~43% for the fly transcriptome). Our approach to estimating the fraction bound assumes that the rank order of the affinities of piRNA-bound PIWI proteins for long RNAs can be approximated by the rank order of the computationally predicted affinities of two naked RNA strands. We therefore assessed the difference in fraction bound between stable and unstable siRNAs by standardizing the fraction bound estimates, i.e., calculating their *Z*-scores. We divided siRNAs by quintile of their fraction remaining in mutants (siRNA abundance in *w*^*1118*^; *hen1*^*f00810*^ divided by siRNA abundance in *w*^*1118*^; +). The *Z*-score of each estimate of fraction bound for unstable siRNAs (≤ 20% of *w^1118^;* +) was then calculated against the background, the median fraction bound for the four other bins (Figure 2B). Consistent with the idea that extensive pairing to long RNAs destabilizes unmethylated siRNAs, the medians of *Z*-scores were >1.96 (i.e., *p* < 0.05) for contiguous pairing to the siRNA seed, central, or 3′ regions (complementary sites starting at nucleotides g2–g5, g10, and g15–g17; Figure 2B). We conclude that both piRNAs and siRNAs are protected from complementarity-dependent destabilization by 3′ terminal 2′-*O*-methylation in flies and likely other arthropods.

### Complementarity-Dependent Destabilization Contributes to Differences in miRNA Decay Rates

Global measurements of small RNA half-lives show that individual miRNA species in the same cell turnover at different rates (Kingston and Bartel, 2019; Reichholf et al., 2019). In TDMD, extensively complementary targets elicit miRNA destruction (Cazalla et al., 2010; Ameres et al., 2010; Xie et al., 2012; Baccarini et al., 2011; Libri et al., 2012; Marcinowski et al., 2012; Rüegger and Großhans, 2012; Lee et al., 2013; de la Mata et al., 2015; Bitetti et al., 2018; Kleaveland et al., 2018; Ghini et al., 2018; Sheu-Gruttadauria et al., 2019a). Do miRNAs that were not documented as TDMD targets but bear abundant complementary sites in the transcriptome also show faster turnover rates?

We used recently reported measurements of fly (Reichholf et al., 2019) and mouse miRNA decay rates (Kingston and Bartel, 2019) to identify highly stable and unstable miRNA species. Again, we used the equilibrium approach to estimate the fraction of stable and unstable miRNAs predicted to bind contiguously complementary sites of various types in the transcriptome (Figure 1A). Supporting the idea that complementarity-dependent destabilization increases miRNA turnover rate, the fraction of miRNA bound to complementary sites was greater for unstable than stable miRNAs. In *Drosophila* S2 cells, mouse embryonic stem cells and contact-inhibited mouse embryonic fibroblasts, pairing of long RNAs to the miRNA central or 3′ regions best explained difference in miRNA turnover rates (Figures 3, 4, and S4).

**Figure 3.**
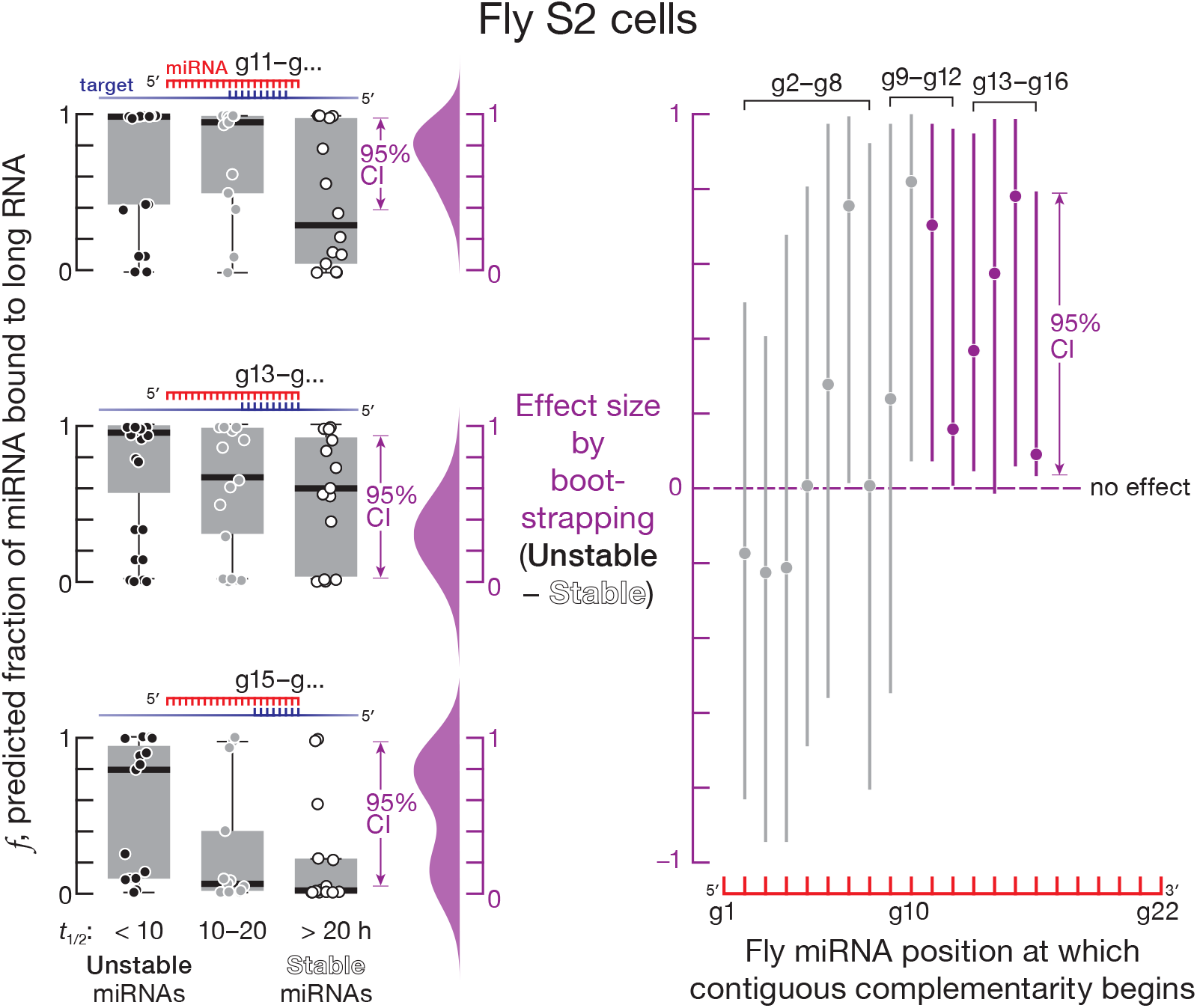
Complementarity-Dependent Destabilization Contributes to Differences in Fly miRNA Half-lives. Left, mean (*n* = 2) predicted fraction of different regions of fly miRNAs bound to complementary sites in the transcriptome. Right, the difference (i.e., unstable miRNAs - stable miRNAs) between the median predicted fraction bound for complementarity starting at miRNA positions g2–g16 for stable (half-lives ≥ 20 hours) and unstable miRNAs (half-lives ≤ 10 hours). Grey: effect size for patterns of complementarity for which the median predicted fraction bound of adjacent quintiles failed to decrease monotonically in the unstable-to-stable direction in Figure S4A. The 95% confidence interval for the effect size of median difference was calculated with 10,000 bootstrapping iterations.

**Figure 4.**
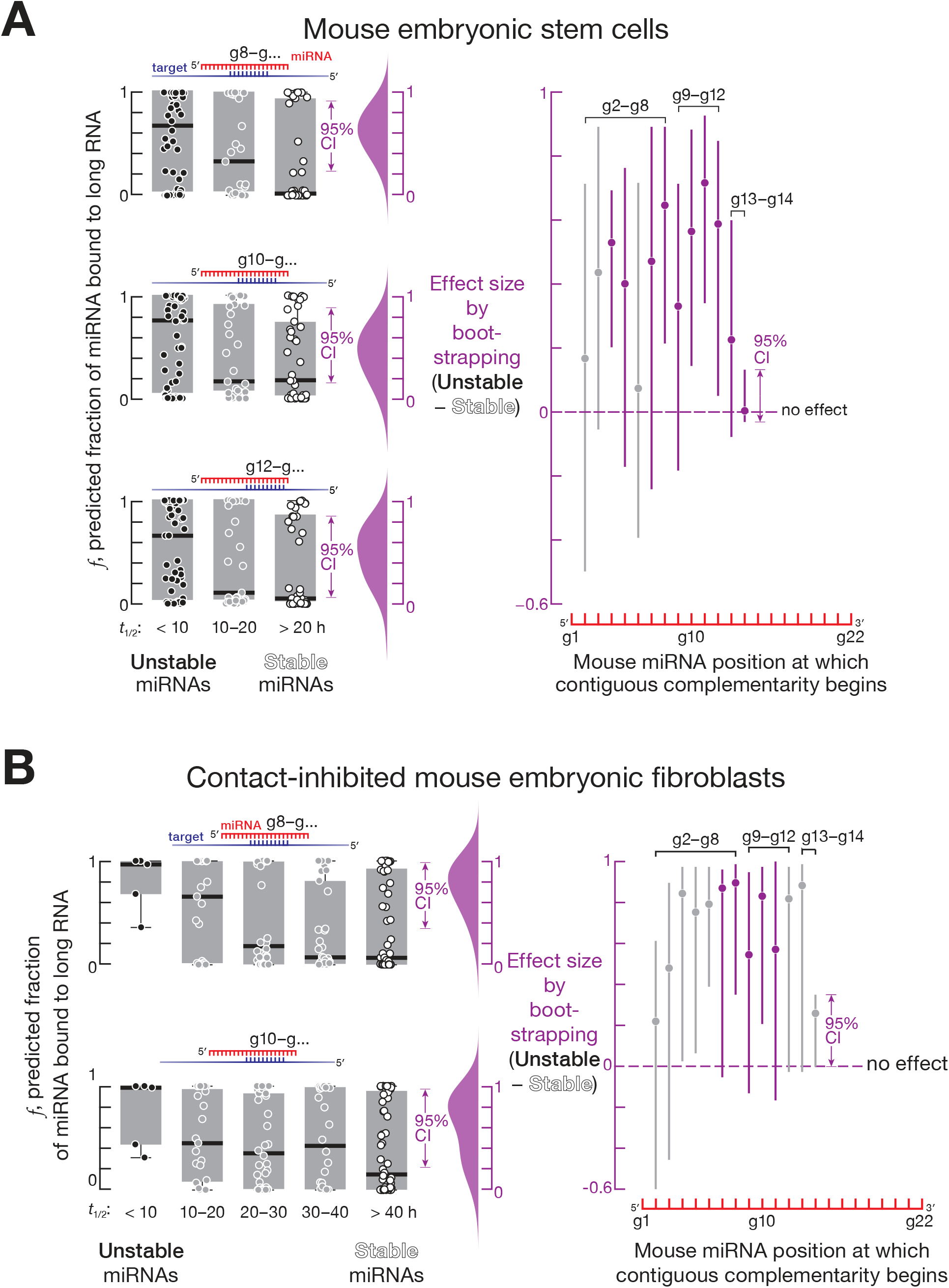
Complementarity-Dependent Destabilization Contributes to Differences in Mouse miRNA Half-lives. (A, B) Left, mean (*n* = 2) predicted fraction of different regions of mouse embryonic stem cell (A) or contact-inhibited mouse embryonic fibroblast (B) miRNAs bound to complementary sites in the transcriptome. Right, the difference (i.e., unstable miRNAs - stable miRNAs) between the median predicted fraction bound for complementarity starting at miRNA positions g2–g14 for stable (half-lives ≥ 20 hours) and unstable miRNAs (half-lives ≤ 10 hours). Grey: effect size for patterns of complementarity for which the median predicted fraction bound of adjacent quintiles failed to decrease monotonically in the unstable-to-stable direction in Figures S4B and S4C. The 95% confidence interval for the effect size of median difference was calculated with 10,000 bootstrapping iterations.

For fly S2 cells, the decay rates of Ago1-bound miRNAs were best explained by contiguous pairing between a long RNA and a miRNA starting at positions g11–g16: e.g., for complementary sites starting at nucleotide g11, the difference in median fraction bound between miRNAs with half-lives < 10 hours and miRNAs with half-lives > 20 hours was ~0.69 (95% CI: 0.06–0.96; Figures 3 and S4A). For mouse embryonic stem cells, contiguous pairing beginning at positions g7–g13 best explained the difference between stable and unstable miRNAs: e.g., for complementary sites starting at miRNA nucleotide g10, the difference in median fraction bound between unstable (half-life < 10 hours) and stable (half-life > 20 hours) miRNAs was ~0.58 (95% CI: 0.15– 0.89; Figures 4A and S4B). For contact-inhibited mouse embryonic fibroblasts, contiguous pairing starting at positions g7–g11 best explained miRNA decay: e.g., for complementary sites starting at miRNA nucleotide g8 the difference in median fraction bound between miRNAs with half-lives < 10 hours and miRNAs with half-lives > 40 hours was ~0.9 (95% CI: 0.35–0.99; Figures 4B and S4C). We did not find evidence for complementarity-dependent destabilization in dividing mouse embryonic fibroblasts (Figure S4D). The differences between animals and cell types may reflect the identity of the Argonaute protein partner of miRNAs or other, yet-to-be-discovered determinants of miRNA stability.

TDMD is triggered by targets that are complementary to both the miRNA seed and miRNA 3′ region but not to ≥ 1 miRNA central nucleotides (Sheu-Gruttadauria et al., 2019a). Our data show that, for mice, complementarity-dependent destabilization of miRNAs is elicited by long RNAs *contiguously* complementary to the miRNA central region (Figures 4A and 4B). For both flies and mice, we also find that, in many cases, complementarity only to the miRNA 3′ region in the absence of a seed match explains differences in miRNA turnover rates whereas pairing to the same 3′ region plus the seed does not: those also containing a seed match as well as contiguous complementarity to miRNA 3′ regions had essentially equivalent differences in fraction bound between stable and unstable miRNAs (Figures 3, 4, S4E, S4F, and S4G; permitting a ≤ 10 nt insertion in the target opposite the central region of miRNA; Sheu-Gruttadauria et al., 2019b; Becker et al., 2019).

Taken together, these data suggest that miRNAs are also subject to complementarity-dependent destabilization and that abundant, high-affinity complementary sites in the transcriptome reduce miRNA half-lives.

### Pre-piRNA Trimming and 2′-*O*-methylation Protect Mouse piRNAs From Different Degradation Mechanisms

In mice, both piRNA 2′-*O*-methylation by HENMT1 and pre-piRNA trimming by PNLDC1 protect piRNAs from degradation (Figure S5A; Lim et al., 2015; Ding et al., 2017; Gainetdinov et al., 2018). Do methylation and trimming protect piRNAs from the same or different degradation mechanisms? Untrimmed pre-piRNAs in *Pnldc1*^*em1Nkn/em1Nkn*^ mice are 2′-*O*-methylated (Nishimura et al., 2018), suggesting that a pathway insensitive to 2′-*O*-methylation degrades pre-piRNAs in the absence of trimming. If different degradation mechanisms act on unmethylated piRNAs and untrimmed pre-piRNAs, then the abundance of the same unmethylated but trimmed piRNA in a *Henmt1* mutant and untrimmed but 2′-*O*-methylated pre-piRNA in a *Pnldc1* mutant are predicted to be uncorrelated. We compared the decrease in abundance of unmethylated piRNAs and the corresponding untrimmed pre-piRNAs by identifying piRNAs in *Henmt1*^*em1/em1*^ and pre-piRNAs in *Pnldc1*^*em1Pdz/em1Pdz*^ (henceforth, *Pnldc1*^*em1/em1*^; Gainetdinov et al., 2018) that began with the same 24-nt sequence (i.e., 5′ prefix). Consistent with the prediction, the decrease of piRNAs in *Henmt1*^*em1/em1*^ and of pre-piRNAs in *Pnldc1*^*em1/em1*^ were poorly correlated (Pearson’s *ρ* = 0.25 and *R*^2^ = 0.06 for spermatogonia; Pearson’s *ρ* = 0.14 and *R*^2^ = 0.02 for primary spermatocytes; Figure 5A). Instead, overlapping but distinct subsets of piRNAs or pre-piRNAs were lost in each mutant, suggesting that 2′-*O*-methylation and trimming protect piRNAs from different degradation mechanisms.

**Figure 5.**
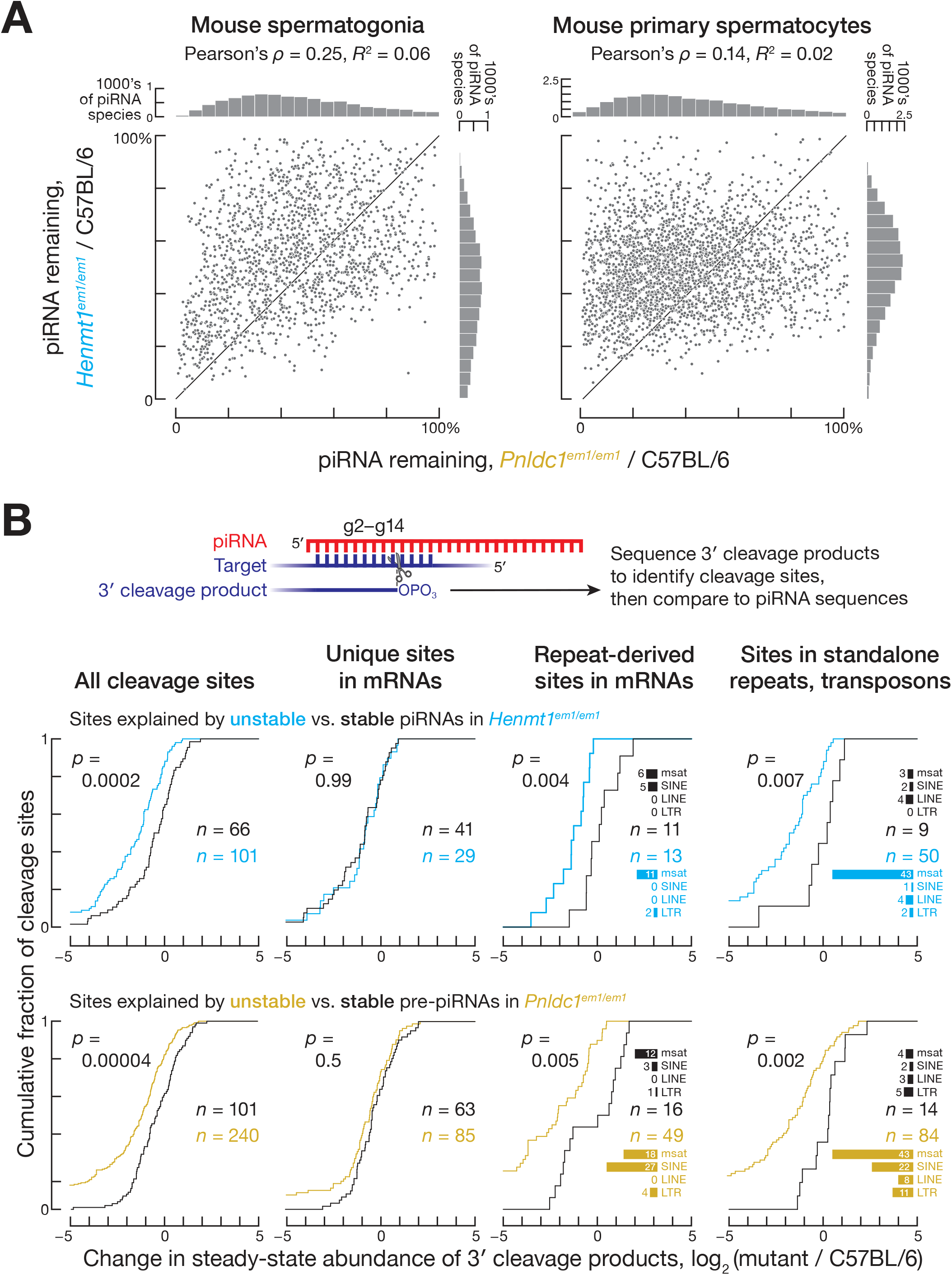
Pre-piRNA Trimming and Methylation Protect Mouse piRNAs From Different Degradation Mechanisms. (A) Mean (*n* = 2) change in mouse piRNA abundance in *Henmt1*^*em1/em1*^ and *Pnldc1*^*em1/em1*^ pre-pachytene piRNAs from FACS-purified spermatogonia (left) and pachytene piRNAs from FACS-purified primary spermatocytes (right). (B) Change in steady-state abundance of 3′ cleavage products explained by contiguous pairing with piRNA nucleotides g2–g14. Data are from FACS-purified primary spermatocytes from *Henmt1*^*em1/em1*^ and *Pnldc1*^*em1/em1*^ mice. Data are from a single representative experiment for piRNAs whose abundance was ≥ 50 molecules per C57BL/6 primary spermatocyte and reduced in both *Henmt1*^*em1/em1*^ and *Pnldc1*^*em1/em1*^ to ≤ 20% of C57BL/6 levels (unstable piRNAs and pre-piRNAs) and to ≥ 80% of C57BL/6 levels (stable piRNAs and pre-piRNAs). *P* values were calculated using two-tailed KS test; msat, microsatellite repeats.

In theory, if the rate of 2′-*O*-methylation were much slower than the rate of destruction of unmethylated, untrimmed pre-piRNAs, then degradation of untrimmed pre-piRNAs could still be explained by complementarity-dependent destabilization. The hypothesis predicts that, like unmethylated piRNAs, unstable, untrimmed pre-piRNAs should have highly abundant, high-affinity complementary sites in the transcriptome. We estimated the fraction of stable (≥ 80% of C57BL/6) and unstable (≤ 20% of C57BL/6) pre-piRNAs bound to complementary sites in transcriptome. Contrary to the prediction, the difference in the median fraction bound for unstable and stable untrimmed pre-piRNAs in *Pnldc1*^*em1/em1*^ (Figures S5B, S5C, S5D, and S5E) was smaller than the corresponding difference for unstable and stable unmethylated piRNAs in *Henmt1*^*em1/em1*^ (Figures 1B, 1C, S1B, and S1C): e.g., in primary spermatocytes for complementarity starting at position g2, the difference in median fraction bound between unstable and stable pre-piRNAs was ~0.21 (95% CI: 0.11– 0.21; Figure S5B) vs. ~0.9 between unstable and stable unmethylated piRNAs (95% CI: 0.87–0.92; Figure 1B). Moreover, in primary spermatocytes, pre-piRNAs were *more* stable when they had highly abundant, high-affinity complementary sites starting at nucleotides g13 to g19 (Figure S5B): for complementarity starting at position g15, the median fraction bound was higher for stable compared to unstable pre-piRNAs (median difference = 0.28; 95% CI: 0.21–0.35; Figure S5B). Thus, the instability of untrimmed pre-piRNAs is unlikely to be driven by complementarity to sequences within long RNAs. We conclude that untrimmed pre-piRNAs and unmethylated piRNAs are degraded by distinct mechanisms.

### Determinants Triggering Degradation of Untrimmed Mouse Pre-piRNAs

Unlike unmethylated piRNAs, the instability of untrimmed pre-piRNAs best correlated with both the identity of the PIWI protein to which the pre-piRNA was bound and the presence of oligoguanine or oligouridine tracts in the pre-piRNA sequence. We sought to compare the stability of untrimmed pre-piRNAs bound to MILI to those bound to MIWI. PIWI proteins and other Argonautes are typically unstable without a small RNA guide (Haase et al., 2010; Zamparini et al., 2011; Derrien et al., 2012; Martinez and Gregory, 2013; Martinez et al., 2013; Smibert et al., 2013; Kobayashi et al., 2019). Because piRNA biogenesis requires binding of PIWI protein to the 5′ end of a pre-pre-piRNA, all pre-piRNAs and piRNAs are anticipated to be bound by PIWI proteins (Gainetdinov et al., 2018). We therefore used the change in the abundance of MIWI and MILI (Figures S6A and S6B) to infer the change in abundance of MIWI- and MILI-bound pre-piRNAs in *Pnldc1*^*em1/em1*^ and piRNAs in *Henmt1*^*em1/em1*^ males. In *Pnldc1*^*em1/em1*^ primary spermatocytes, MIWI abundance was ~30% of C57BL/6, whereas MILI level was ~80% of C57BL/6 (Gainetdinov et al., 2018). In contrast, in *Henmt1*^*em1/em1*^ primary spermatocytes, the abundance of MIWI and MILI declined by similar extents (~70% of C57BL/6 for MIWI and ~80% for MILI; Figure S6B). We conclude that pre-piRNAs bound to MIWI are less stable than those bound to MILI.

Pre-piRNAs bound to MIWI are, on average, ~3 nt longer than their MILI-bound counterparts (Ding et al., 2017; Gainetdinov et al., 2018). Irrespective of the PIWI protein to which they are bound, long pre-piRNAs might be inherently unstable. Our analyses do not support this hypothesis: for MIWI-bound pre-piRNAs, length was not correlated with instability in *Pnldc1*^*em1/em1*^ primary spermatocytes (Spearman’s *ρ* = 0.01; Figure S6C, left). Similarly, the instability of pre-piRNAs bound to MILI in *Pnldc1*^*em1/em1*^ primary spermatocytes did not correlate with the length of the pre-piRNAs bound to MILI (Spearman’s *ρ* = −0.08; Figure S6C, right). We conclude that PIWI protein partner identity, not pre-piRNA length, determines the instability of untrimmed pre-piRNAs.

In addition to PIWI protein identity, the rate of degradation of untrimmed pre-piRNAs also reflected the sequence of the guide RNA itself. In control C57BL/6 primary spermatocytes, the majority of piRNAs bound to MILI are prefixes of piRNA sequences bound to MIWI. Similarly, in *Pnldc1*^*em1/em1*^ primary spermatocytes, the majority of MILI- and MIWI-bound pre-piRNAs share the same 5′ prefix. The stability of such pairs of MILI- and MIWI-bound untrimmed pre-piRNAs was moderately correlated (Pearson’s *ρ* = 0.65, *R*^2^ = 0.42), consistent with a pre-piRNA stability in part reflecting pre-piRNA sequence (Figure S6D). Neither positional mononucleotide (Figure S6E) nor the strength of predicted secondary structures within a pre-piRNA sequence correlated with untrimmed pre-piRNA instability (Spearman’s *ρ* = 0.08; Figure S6F). In contrast, untrimmed pre-piRNA instability correlated with the presence of oligouridine or oligoguanine tracts in the section of pre-piRNA that is removed by trimming (Figure S6G). We observed enrichment of oligouridine or oligoguanine tracts in unstable pre-piRNAs in primary spermatocytes, which express both MILI and MIWI, but not in spermatogonia, which contain only MILI (Figure S6H). These data suggest that some RNA decay machinery recognizes the combination of distal oligouridine or oligoguanine sequences and MIWI itself.

### Tailing and 3′-to-5′ Shortening of Untrimmed pre-piRNAs and Unmethylated piRNAs in Mice

Our analyses suggest that 3′-to-5′ shortening of mature, trimmed piRNAs and 3′ addition of non-templated nucleotides (tailing) to piRNAs or pre-piRNAs play a limited role in the destruction of untrimmed or unmethylated piRNAs. Most unmethylated piRNAs show increased tailing and 3′-to-5′ shortening (Kamminga et al., 2010; Lim et al., 2015; Svendsen et al., 2019; Figures S6I and S6J) irrespective of their stability: unstable unmethylated piRNAs were no more likely to be tailed or shortened than their stable unmethylated brethren (Figures S6J, S6K, and S6L). In *Pnldc1*^*em1/em1*^ primary spermatocytes, tailing of untrimmed pre-piRNAs increased for some but decreased for other species and was not correlated with pre-piRNA instability (Figures S6I, S6K, and S6L). We note that we cannot exclude the possibility that destruction of unmethylated or untrimmed piRNAs requires tailing or 3′-to-5′ shortening, but the rates of such terminal modifications are not rate-determining for piRNA destruction.

### Decreased piRNA Abundance in *Henmt1*^em1/em1^ and *Pnldc1*^em1/em1^ Mouse Spermatocytes Results in Reduced Cleavage of Target mRNAs

Previous studies suggest that pachytene piRNAs regulate their targets by an siRNA-like cleavage mechanism (Reuter et al., 2011; Zhang et al., 2015; Goh et al., 2015; Wu et al., 2020). Consistent with this model, our data show that cleavage is reduced for mRNAs whose slicing is directed by unstable, unmethylated piRNAs or unstable, untrimmed pre-piRNAs in *Henmt1*^*em1/em1*^ and *Pnldc1*^*em1/em1*^ mouse primary spermatocytes.

We sequenced 5′ monophosphorylated long RNAs to identify candidate 3′ cleavage products of piRNA-guided slicing (Figure 5B). To restrict the candidates to high-confidence cleavage sites, we required piRNA nucleotides g2–g14 to pair with the site of complementarity such that the cleavage occurred between target nucleotides t10 and t11 (Reuter et al., 2011; Wang et al., 2014; Zhang et al., 2015; Goh et al., 2015; Wu et al., 2020)). We then classified the putative 3′ cleavage products by the stability of the piRNAs in *Henmt1*^*em1/em1*^ or pre-piRNAs *Pnldc1*^*em1/em1*^ predicted to generate them. We find that in *Henmt1*^*em1/em1*^ primary spermatocytes, the abundance of 3′ cleavage products produced by unstable piRNAs decreased more than those produced by stable piRNAs (two-tailed KS test, *p* = 0.0002; Figure 5B). Similarly, in *Pnldc1*^*em1/em1*^ primary spermatocytes, the abundance of 3′ cleavage products generated by unstable pre-piRNAs decreased more than those generated by stable pre-piRNAs (two-tailed KS test, *p* = 0.00004; Figure 5B).

For both *Henmt1*^*em1/em1*^ and *Pnldc1*^*em1/em1*^ mutants, piRNA-directed, 3′ cleavage products mapped to both mRNAs and solitary transposon and repeat insertions (Figure 5B). Many cleavage sites in mRNAs corresponded to transposon- or repeat-derived sequences (Figure 5B). In fact, the decrease in 3′ cleavage product abundance in *Henmt1*^*em1/em1*^ and *Pnldc1*^*em1/em1*^ mutants was greater for these sites than for 3′ cleavage sites mapping to unique portions of mRNAs (Figure 5B).

The majority of repeat-derived cleavage sites were in microsatellite repeats and muroid-specific SINE transposons (Figure 5B). Transposon subfamily age has been estimated using the relative extent to which one transposon subfamily has integrated into another (Giordano et al., 2007). Using this information, we find that ~85% of LINE- and LTR transposon-derived cleavage products mapped to evolutionarily older subfamilies of these transposon classes (16 of 19 LINE and 21 of 25 LTR transposons). Yet both evolutionarily younger and older subfamilies of LINE and LTR elements were derepressed in *Henmt1*^*em1/em1*^ and *Pnldc1*^*em1/em1*^ primary spermatocytes (Table S1, Figures S7A and S7B; Giordano et al., 2007).

In *Henmt1*^*em1/em1*^, the decreased abundance of individual unmethylated piRNAs explained the increased steady-state level of 15 mRNAs. We note that the decrease in unmethylated piRNA abundance was first detected in primary spermatocytes, but increased target RNA abundance lagged and was often observed only in secondary spermatocytes or round spermatids (Table S2). This phenomenon was observed previously for mice mutant for a piRNA-producing locus on chromosome 6 (Wu et al., 2020).

The reduction in abundance of untrimmed pre-piRNAs in *Pnldc1*^*em1/em1*^ primary spermatocytes similarly explained the increase of steady-state levels of 26 mRNAs observed in *Pnldc1*^*em1/em1*^ mutant primary spermatocytes, secondary spermatocytes, or round spermatids (Table S2). The cleavage sites in many of these mRNAs map to transposon- or repeat-derived sequences. Consistent with the finding that different subsets of piRNAs are unstable in *Henmt1*^*em1/em1*^ and *Pnldc1*^*em1/em1*^animals, just three mRNAs targeted by unstable piRNAs or pre-piRNAs were common to the two mutants (Table S2).

### The Mouse piRNA Pathway Collapses in the Absence of Both Trimming and 2′-O-Methylation

Single mutant *Henmt1*^*em1/em1*^ and *Pnldc1*^*em1/em1*^ spermatogonia exhibited a ~2–3-fold decline in piRNA abundance (Figures 6A and 6B). Trimming and 2′-O-methylation protect overlapping but distinct sets of piRNAs (Figure 5A), so removing both PNLDC1 and HENMT1 should cause a greater decrease in piRNA abundance. Consistent with the prediction, the piRNA pathway collapsed in the spermatogonia of *Henmt1*^*em1/em1*^; *Pnldc1*^*em1/em1*^ double mutants: piRNA abundance decreased ~sixfold (Figures 6A and 6B). Consistent with the larger loss of piRNAs, double mutant spermatogonia displayed a more severe phenotype than *Henmt1*^*em1/em1*^ or *Pnldc1*^*em1/em1*^ mice: germ cells in double mutants developed no further than the pachytene stage of meiosis, whereas the single mutants arrest after concluding meiosis (Figures 6C, 6D, and 6E).

**Figure 6.**
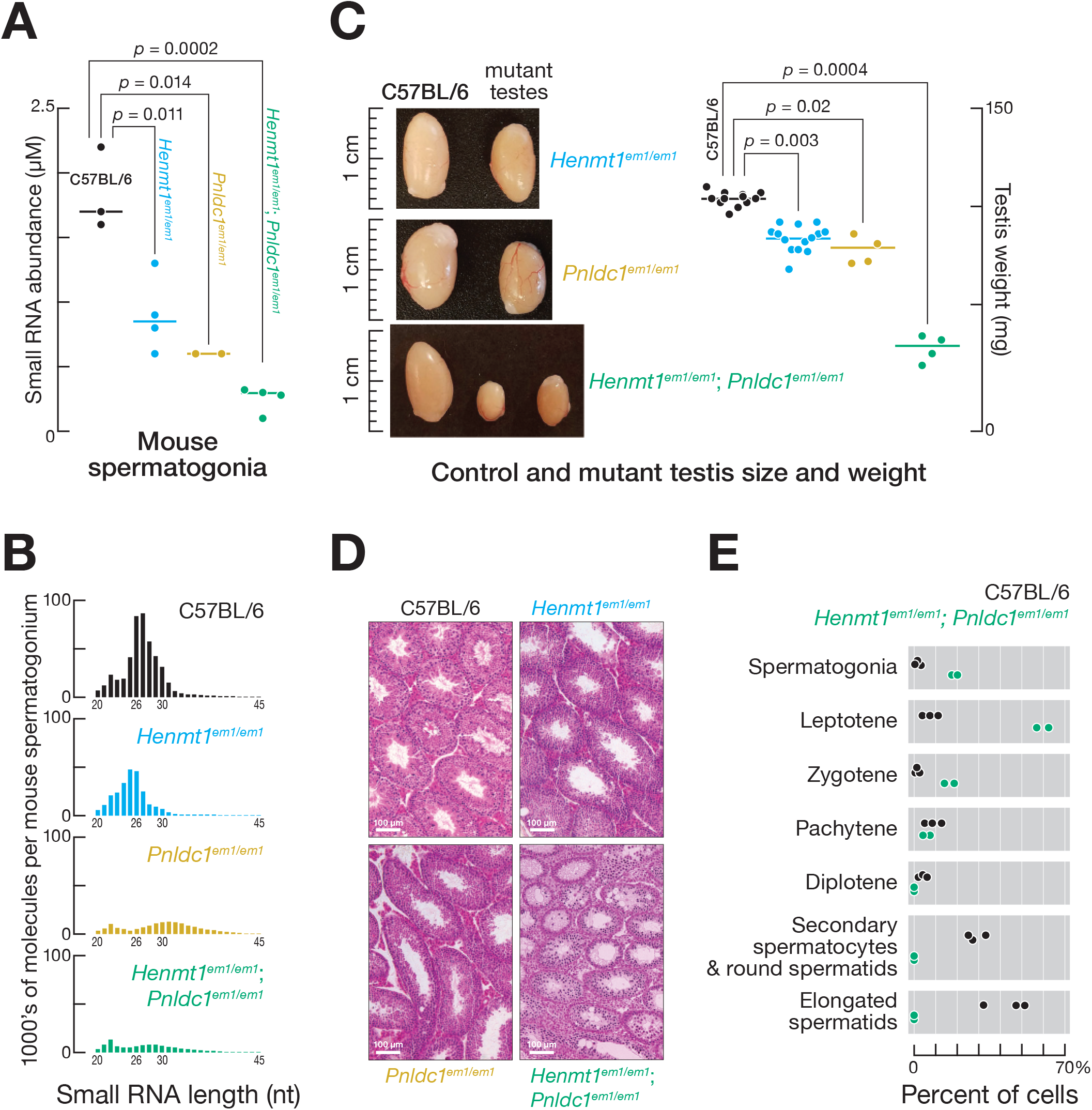
piRNA Pathway Collapses in *Henmt1*^*em1/em1*^; *Pnldc1*^*em1/em1*^ double mutants. (A) Median (*n* = 2–4) abundance of mouse pre-pachytene piRNAs in FACS-purified spermatogonia (24–33-nt small RNAs for C57BL/6 and *Henmt1*^*em1/em1*^; 24–45-nt small RNAs for *Pnldc1*^*em1/em1*^ and *Pnldc1*^*em1/em1*^; *Henmt1*^*em1/em1*^). (B) Mean (*n* = 2) small RNA length profiles for ≥ 20-nt small RNAs from C57BL/6, *Henmt1*^*em1/em1*^, *Pnldc1*^*em1/em1*^ and *Henmt1*^*em1/em1*^; *Pnldc1*^*em1/em1*^ FACS-purified spermatogonia. (C) Size and median (*n* = 4–13) weight of testes from 2–4 month-old C57BL/6, *Henmt1*^*em1/em1*^, *Pnldc1*^*em1/em1*^ and *Henmt1*^*em1/em1*^; *Pnldc1*^*em1/em1*^ mice. *P* values calculated using Mann–Whitney *U* test. (D) Hematoxylin and eosin staining of sections from 2–4 month-old C57BL/6, *Henmt1*^*em1/em1*^, *Pnldc1*^*em1/em1*^and *Henmt1*^*em1/em1*^; *Pnldc1*^*em1/em1*^ testes. (E) Germ cell type composition of C57BL/6 and *Henmt1*^*em1/em1*^; *Pnldc1*^*em1/em1*^ testes. Each data point corresponds to one animal.

Unlike in *Henmt1*^*em1/em1*^ or *Pnldc1*^*em1/em1*^ single mutant spermatogonia, steady-state abundance of transposon mRNAs increased in *Henmt1*^*em1/em1*^; *Pnldc1*^*em1/em1*^ spermatogonia (~2-fold for L1-A, *p*_adj_ = 10^−8^; ~1.7-fold for L1-Gf, *p*_adj_ = 0.045; ~3.6-fold for IAPEY4, *p*_adj_ = 2 × 10^−8^; Figure S7C, Table S1). piRNAs bound to MIWI2 direct DNA methylation of transposons in the fetal testis (Aravin et al., 2008; Kuramochi-Miyagawa et al., 2008). We used targeted bisulfite sequencing to measure the extent of DNA methylation of evolutionarily younger LINE subfamilies L1-Gf and L1-A, and IAP LTR elements. Two pairs of primers specific to thousands of L1-Gf and L1-A genomic copies and a primer pair specific to a single copy of IAP LTR element (Kojima-Kita et al., 2016; Nishimura et al., 2018) were used to amplify bisulfite treated DNA. In both C57BL/6 and *Henmt1*^*em1/em1*^; *Pnldc1*^*em1/em1*^ spermatogonia, the median level of CpG methylation was ≥ 80% (Figure S7D), suggesting that transposon derepression in the double mutants reflects impaired post-transcriptional silencing.

Our data suggest that, in the double mutant, pre-piRNAs are degraded both by complementarity-dependent destabilization acting on their unmethylated 3′ termini and by a pathway specifically targeting untrimmed pre-piRNAs. The decreased abundance of untrimmed, unmethylated pre-piRNAs in double mutant spermatogonia did not correlate strongly with the decrease of unmethylated piRNAs in *Henmt1*^*em1/em1*^ (Pearson’s *ρ* = 0.33, *R*^2^ = 0.11; Figure 7A, right) or the decrease in untrimmed pre-piRNAs in *Pnldc1*^*em1/em1*^ single mutant spermatogonia (Pearson’s *ρ* = 0.63, *R*^2^ = 0.40; Figure 7A, center). These data suggest that unmethylated, untrimmed pre-piRNAs with abundant, high-affinity complementary sites in transcriptome are unstable.

**Figure 7.**
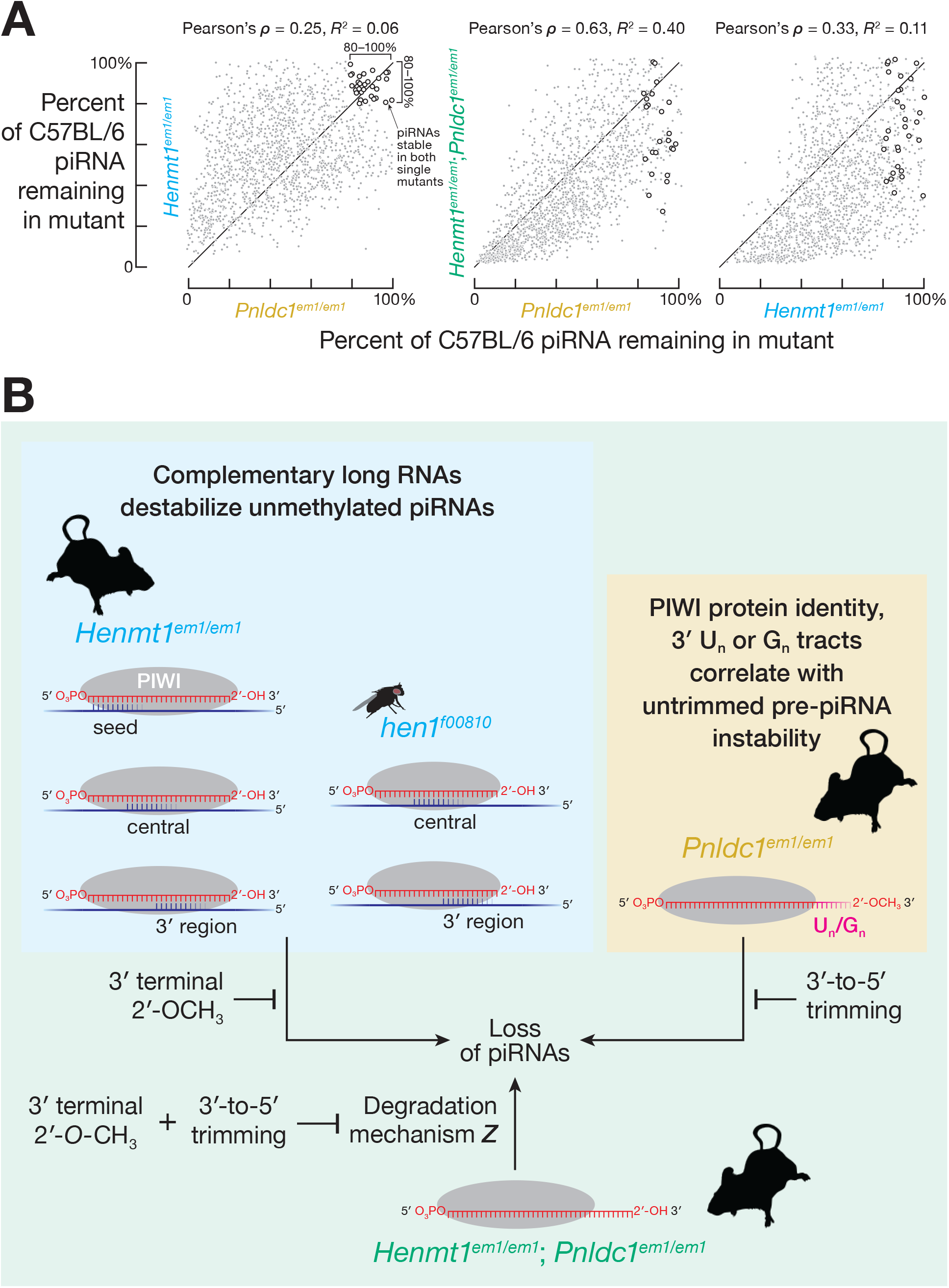
Three Distinct Pathways Destroy Unmethylated, Untrimmed Pre-piRNAs. (A) Mean (*n* = 2) change in piRNA abundance for FACS-purified spermatogonia from *Henmt1*^*em1/em1*^ and *Pnldc1*^*em1/em1*^ single mutants and *Henmt1*^*em1/em1*^; *Pnldc1*^*em1/em1*^ double mutants. Open circles indicate piRNAs whose abundance in both *Henmt1*^*em1/em1*^ and *Pnldc1*^*em1/em1*^ single mutants remained ≥ 80% of C57BL/6 levels. (B) A model for how trimming and 2′-*O*-methylation collaborate to stabilize mouse piRNAs.

To test this prediction, we compared the fraction of stable (≥ 80% of C57BL/6 levels) and unstable (≤ 20% of C57BL/6 levels) pre-piRNAs bound to complementary sites in *Henmt1*^*em1/em1*^; *Pnldc1*^*em1/em1*^ double mutant spermatogonia. Consistent with complementarity-dependent degradation of unmethylated pre-piRNAs in *Henmt1*^*em1/em1*^; *Pnldc1*^*em1/em1*^ spermatogonia, the fraction bound to long RNAs was higher for unstable compared to stable pre-piRNAs: e.g., for complementary sites starting at nucleotide g2, the difference in the median fraction between stable and unstable pre-piRNAs was ~0.62 (95% CI: 0.56–0.68; Figures S7E and S7F).

We also find evidence for a third degradation pathway destroying untrimmed, unmethylated pre-piRNAs in *Henmt1*^*em1/em1*^; *Pnldc1*^*em1/em1*^ double mutants. If piRNA degradation in double mutants reflects the joint action of only two degradation pathways, piRNAs that are stable in both *Henmt1*^*em1/em1*^and *Pnldc1*^*em1/em1*^ single mutants should be stable in the double mutants. Yet of the ~220 piRNAs that were stable in both *Henmt1*^*em1/em1*^ and *Pnldc1*^*em1/em1*^ single mutants, just half remained stable in the *Henmt1*^*em1/em1*^; *Pnldc1*^*em1/em1*^ double mutants (Figure 7A). These data suggest that untrimmed, unmethylated pre-piRNAs in *Henmt1*^*em1/em1*^; *Pnldc1*^*em1/em1*^ are degraded by a pathway that can act on neither trimmed, unmethylated piRNAs in *Henmt1*^*em1/em1*^ nor untrimmed, methylated pre-piRNAs in *Pnldc1*^*em1/em1*^ (Figure 7B).

## DISCUSSION

The data presented here show that 3′-to-5′ trimming by PNLDC1 and 2′-*O*-methylation by HENMT1 protect mouse piRNAs from two separate degradation mechanisms, and in the absence of both maturation steps, untrimmed, unmethylated pre-piRNAs are destabilized by the two destruction pathways as well as an additional third mechanism that cannot act on trimmed but unmethylated piRNAs or untrimmed but methylated pre-piRNAs (Figure 7B).

In mice, 2′-*O*-methylation protects piRNAs from decay triggered by binding to extensively complementary RNAs. In mammals, the testis has a highly complex transcriptome, with as many as ~27,300 distinct mRNAs and lncRNAs (for comparison, liver tissues express ~16,500 different transcripts; Soumillon et al., 2013). Consequently mouse piRNAs have a greater probability of encountering a complementary target than small RNAs in the soma.

Complementarity-dependent destabilization of piRNAs is conserved in animals as evolutionarily distant as mice and flies, whose last common ancestor existed ~800 million years ago (Kumar et al., 2017). For piRNAs in mice and flies, complementarity to different regions of the guide trigger destabilization. We speculate that this distinction may be partly attributed to the different PIWI protein partners of piRNAs. In flies, 2′-*O*-methylation also protects siRNAs from complementarity-dependent destabilization. Our data, together with studies in Cnidaria, Ciliophora, and plants (Park et al., 2002; Chen et al., 2002; Yu et al., 2005; Li et al., 2005; Kurth and Mochizuki, 2009), suggest an ancestral function of 2′-*O*-methylation in protecting small silencing RNAs from complementarity-dependent destabilization.

If 2′-*O*-methylation protects piRNAs and siRNAs from nucleases, the protection mechanism is unlikely to be explained by the two-to-six-fold greater affinity of the PAZ domain of PIWI proteins to a 2′-*O*-methylated compared to an unmethylated guide (Tian, 2011; Simon et al., 2011). To the contrary: we propose that the need to protect piRNAs by 3′ terminal 2′-*O*-methylation put pressure on PIWI proteins to accommodate the 2′-*O*-methyl moiety in their PAZ domain. This hypothesis predicts that in a mouse expressing a PIWI protein with a mutated PAZ domain, piRNA stability should not be impacted.

miRNAs are not 2′-*O*-methylated in most animals and were likely under evolutionary pressure to avoid extensive pairing with transcripts (Ameres et al., 2010). Supporting this view, we show that miRNAs are subject to complementarity-dependent destabilization, and that miRNA decay rates are, in part, determined by the abundance and affinity of their complementary sites. Unlike TDMD (Sheu-Gruttadauria et al., 2019a), complementarity-dependent destabilization of mouse miRNAs can be triggered by long RNAs bearing contiguous pairing to the miRNA central region and that complementarity-dependent destabilization of mouse and fly miRNAs can be elicited by contiguous pairing to the miRNA 3′ region in the absence of seed pairing. The exact molecular mechanism of complementarity-dependent destabilization and whether TDMD and complementarity-dependent destabilization of small RNAs are overlapping or distinct molecular pathways remains to be assessed (De et al., 2013; Park et al., 2017; Kleaveland et al., 2018; Sheu-Gruttadauria et al., 2019a).

The susceptibility of untrimmed pre-piRNAs to degradation does not depend on complementarity to long RNAs, but is determined by both loading into MIWI rather than MILI and the presence of oligouridine or oligoguanine tracts in the trimmed portion of the pre-piRNA. Pre-piRNA trimming by PNLDC1 is also required for stabilizing mature piRNAs in silkmoth (Izumi et al., 2016; Izumi et al., 2020). PNLDC1 is present in most animals, except fish and dipteran insects, whose pre-piRNAs are just a few nucleotides longer than mature piRNAs (Hayashi et al., 2016; Gainetdinov et al., 2018). Similarly, lengthening of miRNAs by viral poly(A) polymerase results in their destabilization (Backes et al., 2012), suggesting that, in general, small RNAs are produced at or mature to an optimal length that enhances their stability. In flies, 3′ terminal trimming by the 3′-to-5′ exonuclease Nibbler is required for the biogenesis of piRNAs loaded in cytoplasmic PIWI proteins (Hayashi et al., 2016). It is not known why worm piRNAs (21U-RNAs) are stable when untrimmed (Tang et al., 2016).

The finding that pre-piRNA trimming and 2′-*O*-methylation act additively to protect different subsets of piRNAs from distinct decay mechanisms offers an explanation for the surprisingly mild phenotype—post-meiotic spermatogenic arrest— of *Henmt1*^*em1/em1*^ and *Pnldc1*^*em1/em1*^single mutants (Lim et al., 2015; Zhang et al., 2017; Ding et al., 2017; Nishimura et al., 2018): removing both PNLDC1 and HENMT1 results in the collapse of the piRNA pathway and the arrest of spermatogenesis at the onset of meiosis as observed for mice deficient for other piRNA biogenesis proteins (Tanaka et al., 2000; Kuramochi-Miyagawa et al., 2004; Carmell et al., 2007; Soper et al., 2008; Ma et al., 2009; Shoji et al., 2009; Yoshimura et al., 2009; Zheng et al., 2010; Frost et al., 2010; Huang et al., 2011; Watanabe et al., 2011). By collaborating, 3′-to-5′ trimming and 2′-*O*-methylation maintain the high steady-state abundance required for the piRNA pathway to function.

## ACCESSION NUMBERS

Sequencing data are available from the National Center for Biotechnology Information Small Read Archive using accession number PRJNA660633.

## ACKNOWLEDGEMENTS

We thank UMass FACS Core for help sorting mouse germ cells; the UMass Transgenic Animal Modeling Core for help generating *Pnldc1*^*em1Pdz/em1Pdz*^ and *Henmt1*^*em1Pdz/em1Pdz*^ mice; members of the Zamore and Mello laboratory for discussions and critical comments on the manuscript; Dimas Echeverria Moreno, Matthew R Hassler and Jacquelyn Sousa from Khvorova laboratory for the technical assistance. This work was supported in part by National Institutes of Health grants GM65236 and P01HD078253 to P.D.Z.

## AUTHOR CONTRIBUTIONS

I.G., C.C, K.J. and P.D.Z. conceived and designed the experiments. C.C., I.G., K.C., P.A., J.V.B., and D.M.O. performed the experiments. I.G. analyzed the sequencing data. I.G., C.C., and P.D.Z. wrote the manuscript.

## SUPPLEMENTAL FIGURE TITLES AND LEGENDS

**Figure S1.**
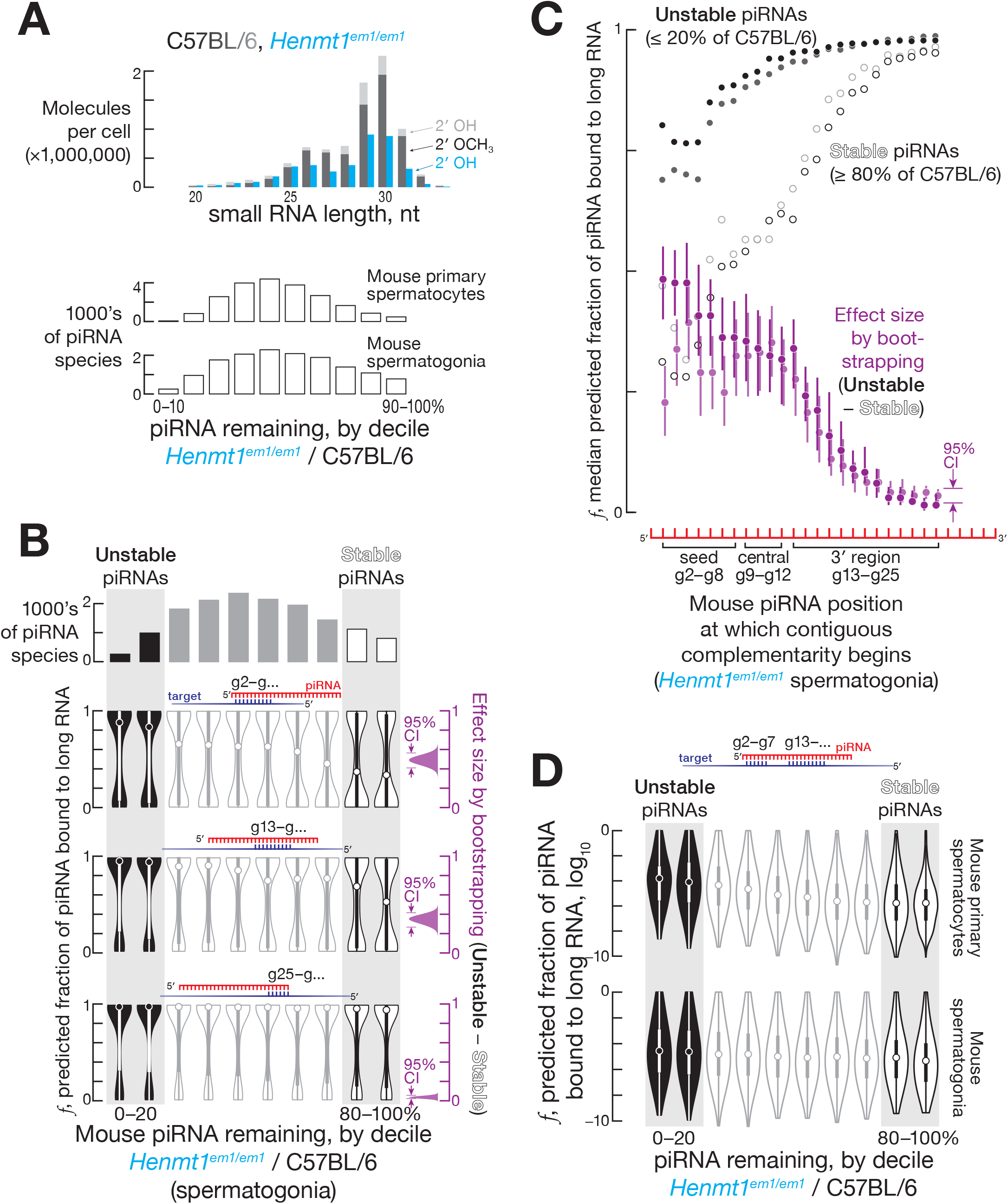
**Related to Figure 1** (A) Mean (*n* = 2) length profiles of ≥ 20-nt small RNAs from *Henmt1*^+/*em1*^ and *Henmt1^em1/^*^em1^ FACS-purified primary spermatocytes. (B) Predicted fraction bound to complementary sites in the transcriptome for different regions of mouse pre-pachytene piRNAs from FACS-purified *Henmt1*^*em1/em1*^ spermatogonia. The 95% confidence interval for the effect size of median difference was calculated with 10,000 bootstrapping iterations. (C) Median predicted fraction bound of spermatogonial pre-pachytene piRNAs for complementarity starting at piRNA positions g2–g25 for stable (≥ 80% of C57BL/6 levels in *Henmt1*^*em1/em1*^) and unstable piRNAs (≤ 20% of C57BL/6 levels in *Henmt1*^*em1/em1*^), as well as the difference between the two (i.e., unstable piRNAs − stable piRNAs). Two independent experiments are shown in different shades of the same color. The 95% confidence interval for the effect size of median difference was calculated with 10,000 bootstrapping iterations. (D) Predicted fraction of pachytene piRNAs from FACS-purified mouse primary spermatocytes bound to sites in the transcriptome bearing g2–g7 seed matches as well as complementarity starting at piRNA nucleotide g13.

**Figure S2.**
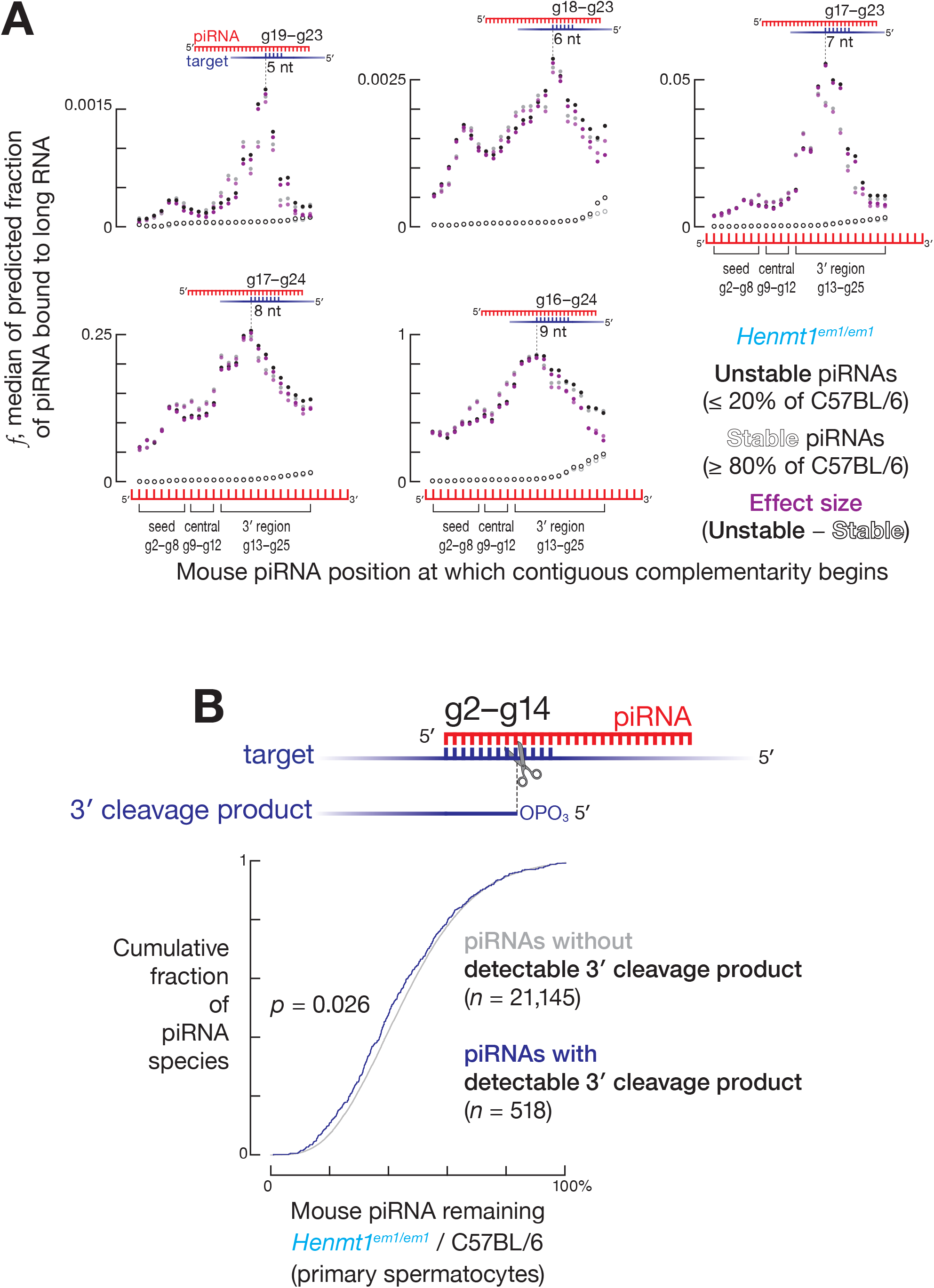
**Related to Figure 1** (A) Predicted fraction bound for piRNAs from FACS-purified mouse primary spermatocytes for complementarity starting at positions g2–g25 for stable (≥ 80% of C57BL/6 levels in *Henmt1*^*em1/em1*^) and unstable piRNAs (≤ 20% of C57BL/6 levels in *Henmt1*^*em1/em1*^), as well as the difference between the two (i.e., unstable piRNAs - stable piRNAs). Data are shown separately for 5-nt, 6-nt, 7-nt, 8-nt, and 9-nt stretches of complementarity. Two independent experiments are shown in different shades of the same color. The 95% confidence interval for the effect size of median difference was calculated with 10,000 bootstrapping iterations. (B) Change in abundance of piRNAs from *Henmt1*^*em1/em1*^ FACS-purified, mouse primary spermatocytes for those piRNAs with and without detectable 3′ cleavage products. *P* value calculated using two-tailed KS test.

**Figure S3.**
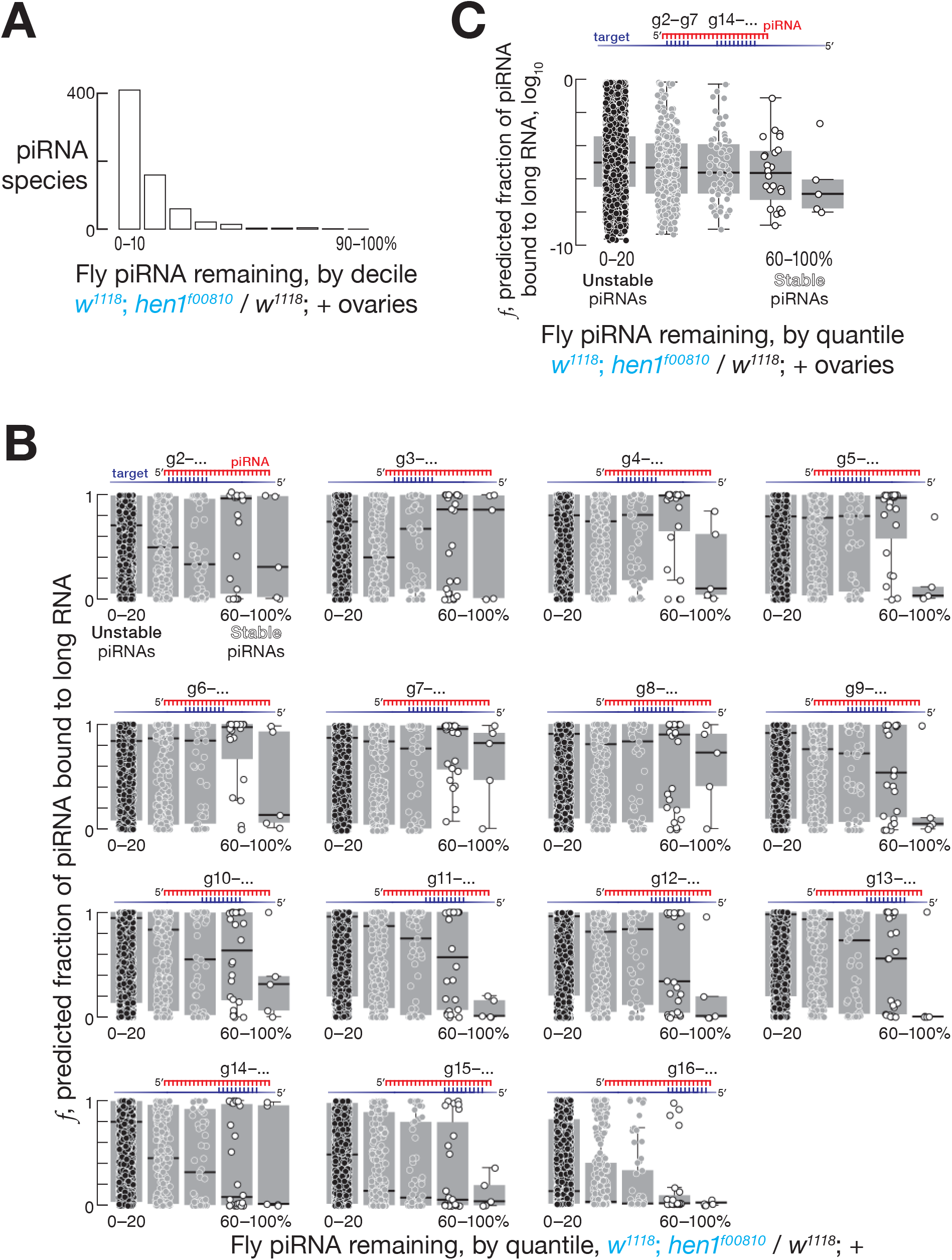
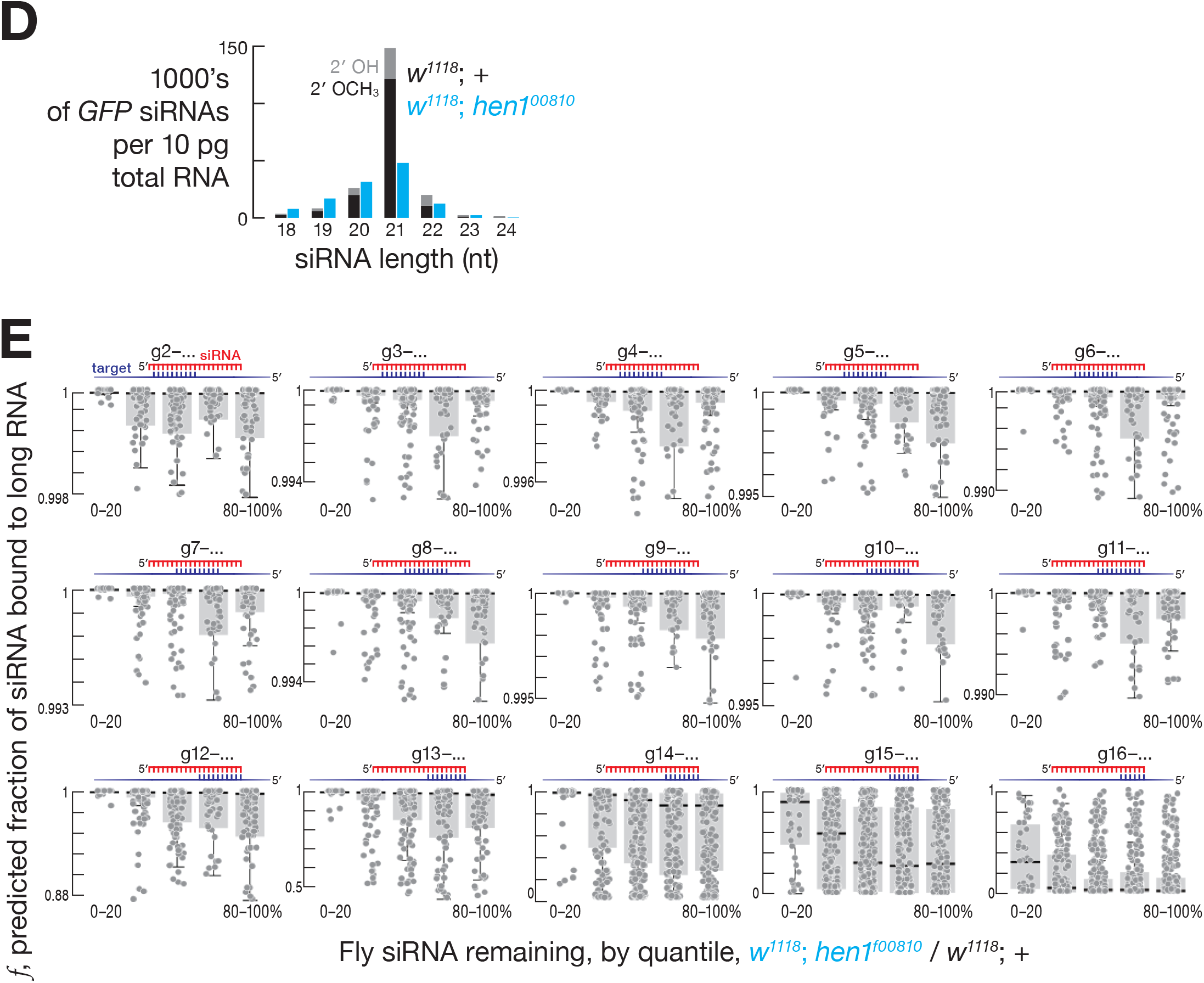
**Related to Figure 2** (A) Change in abundance of *Drosophila melanogaster* piRNAs between control (*w*^*1118*^) and *hen1*^*f00810*^ ovaries. Data represent the mean of two independent experiments. (B) Predicted fraction of different regions of fly piRNAs bound to complementary sites in the transcriptome. Data are for piRNAs in fly ovaries. Data represent the mean of two independent experiments. (C) For piRNAs in fly ovaries bearing g2–g7 seed pairing sites, predicted fraction of piRNAs bound to complementary sites in the transcriptome starting at piRNA nucleotide g14. Data represent the mean of two independent experiments. (D) Mean (*n* = 2) length profiles of GFP-derived siRNAs in control (*w*^*1118*^) and *hen1*^*f00810*^ flies. (E) Fraction of fly siRNAs bound to sites in the transcriptome complementary to different siRNA regions. Data are for GFP-derived siRNAs in heads from *hen1*^*f00810*^ flies.

**Figure S4.**
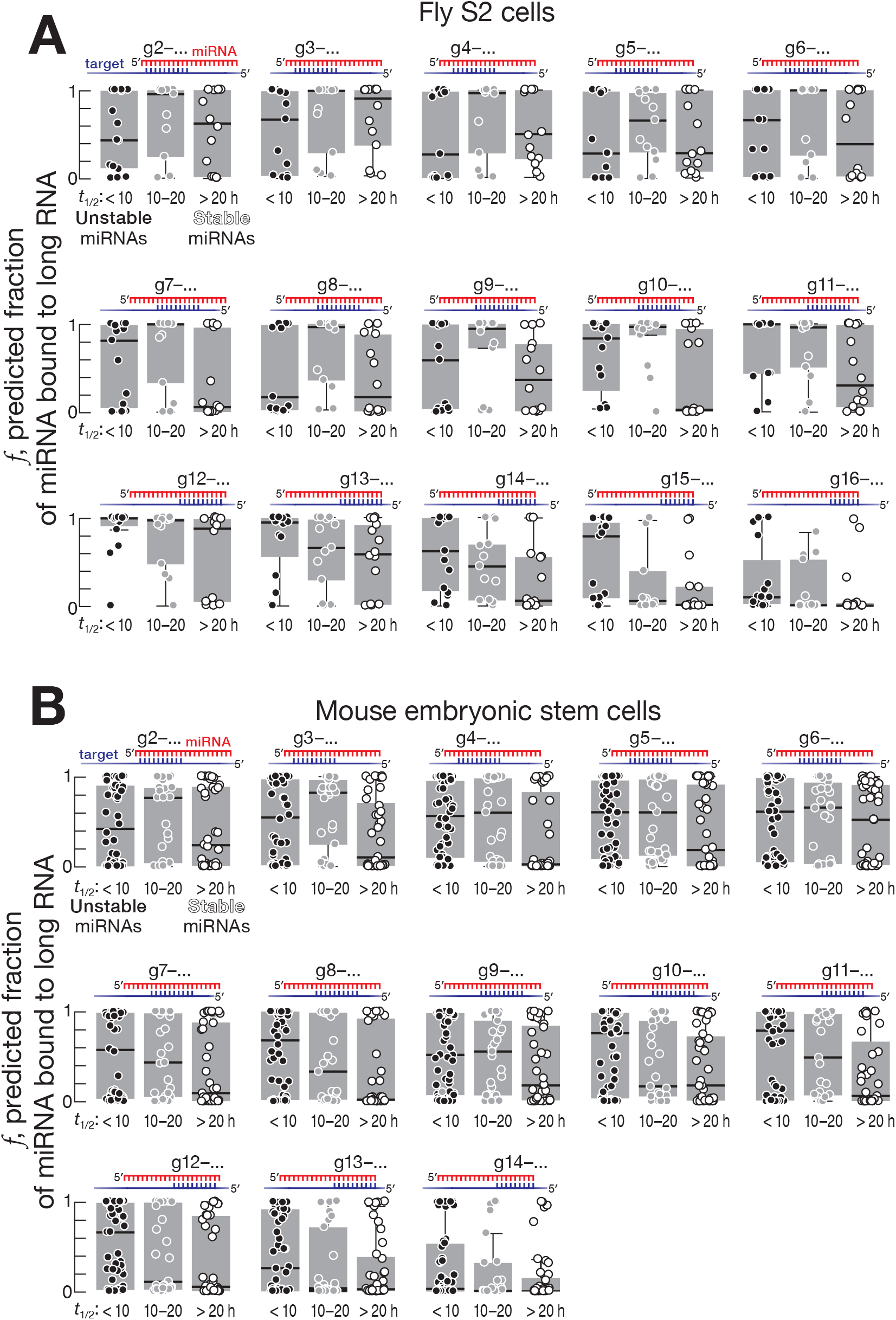
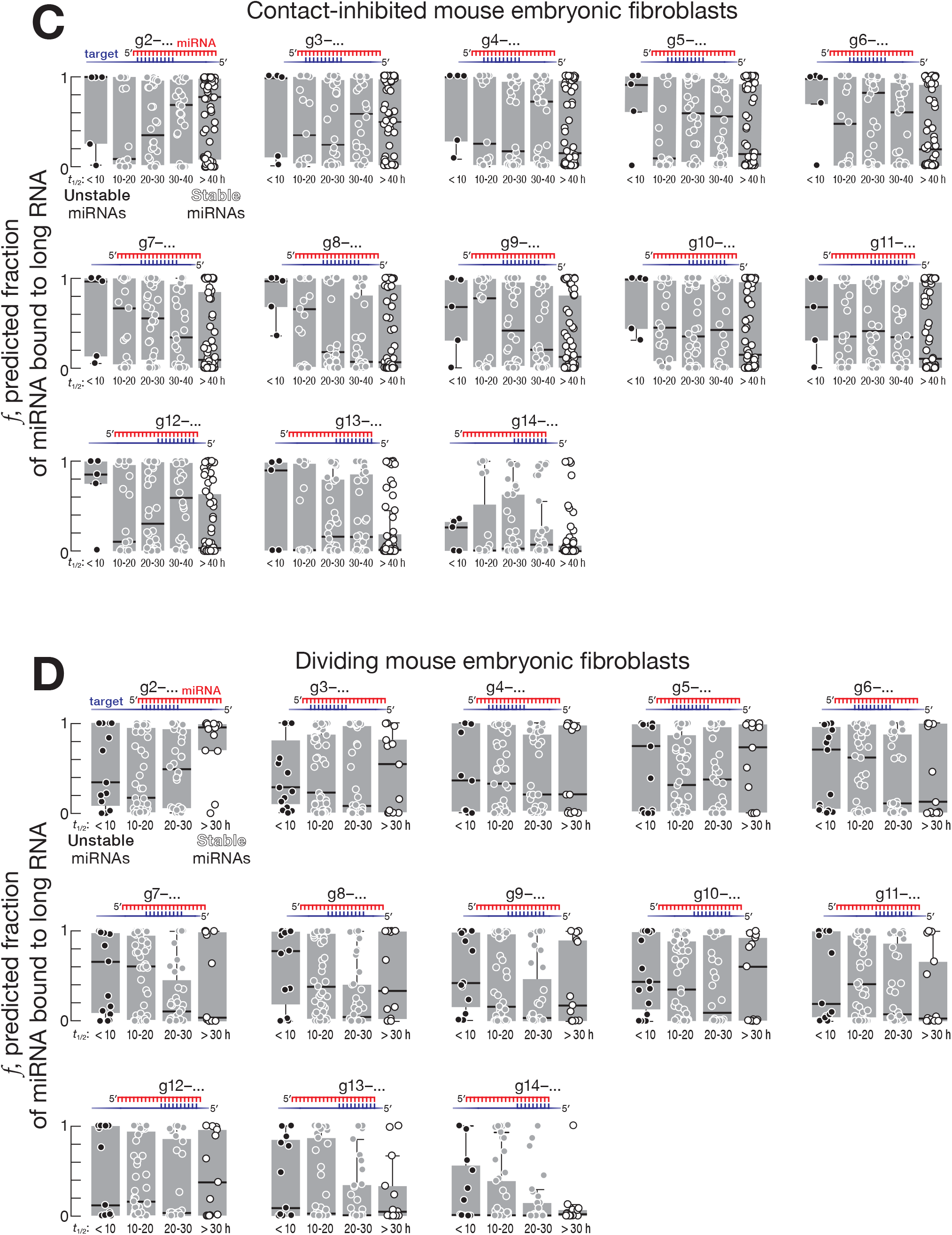
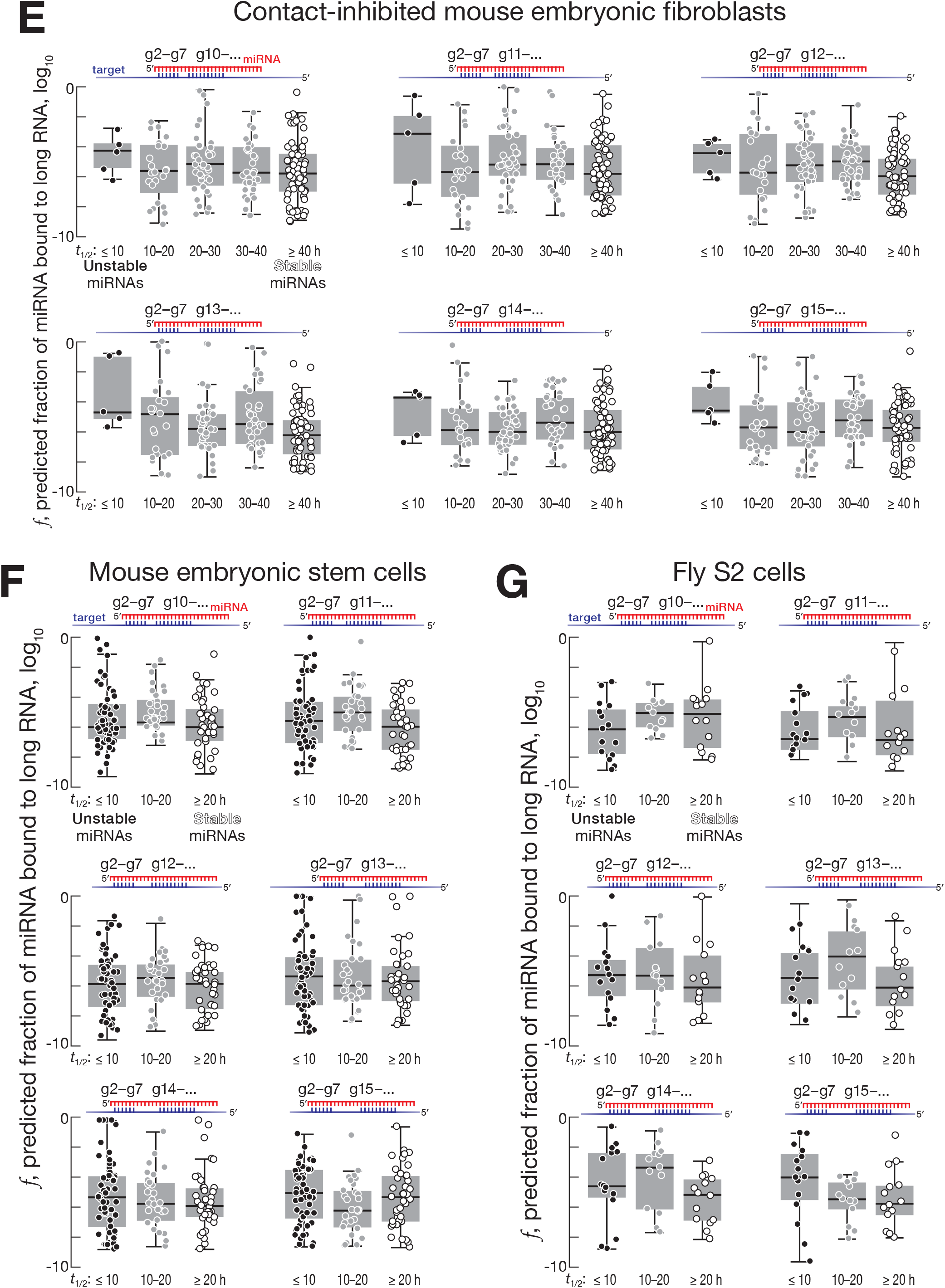
**Related to Figures 3 and 4.** (A, B, C, D) The fraction of miRNAs bound to sites in the transcriptome complementary to different miRNA regions for (A) fly S2 cells, (B) mouse embryonic stem cells, (C) contact-inhibited mouse embryonic fibroblasts, and (D) dividing mouse embryonic fibroblasts. (E, F, G) The same analysis as in (A)–(D) but for miRNAs also bearing g2–g7 seed complementarity to sites in the transcriptome.

**Figure S5.**
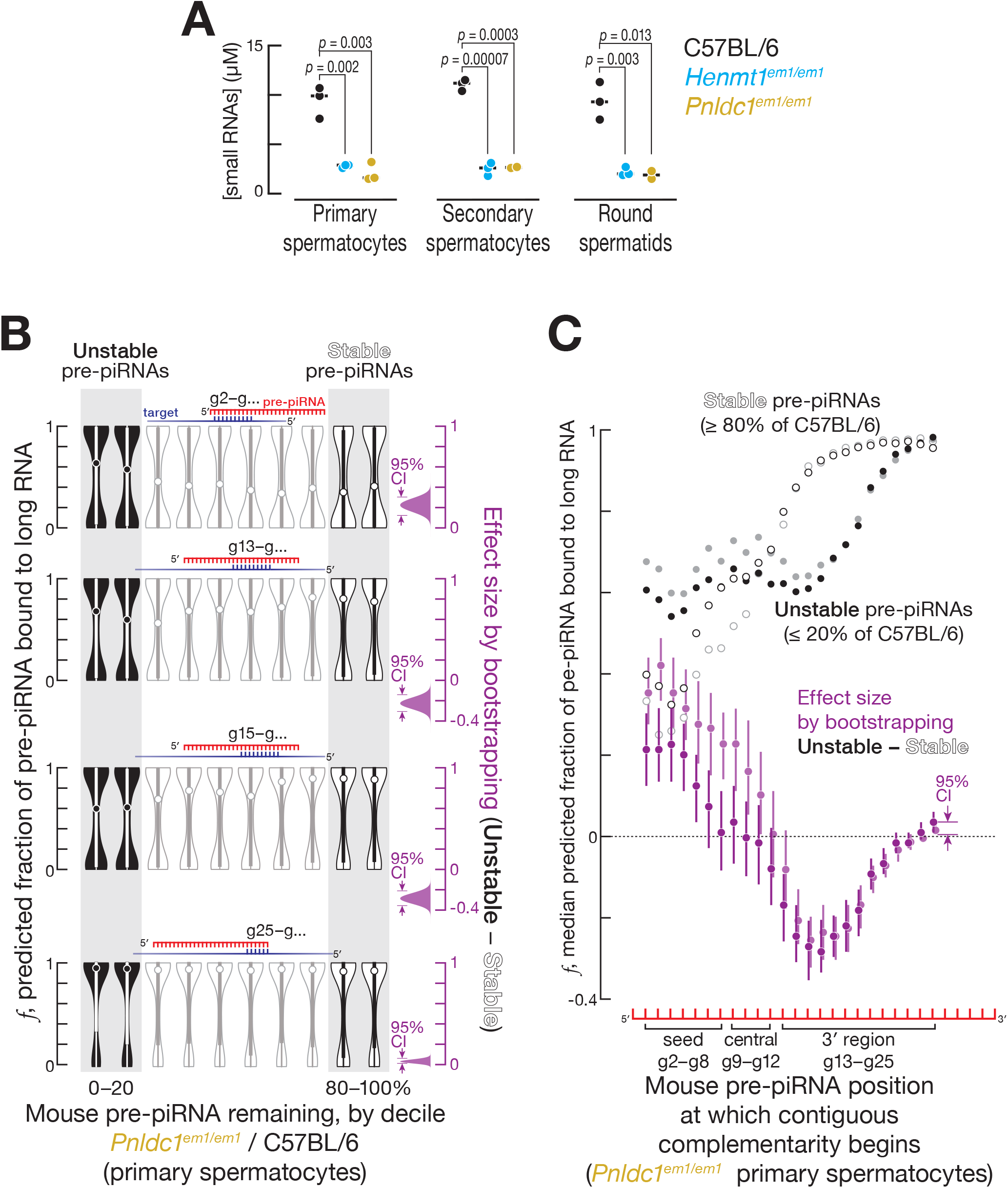
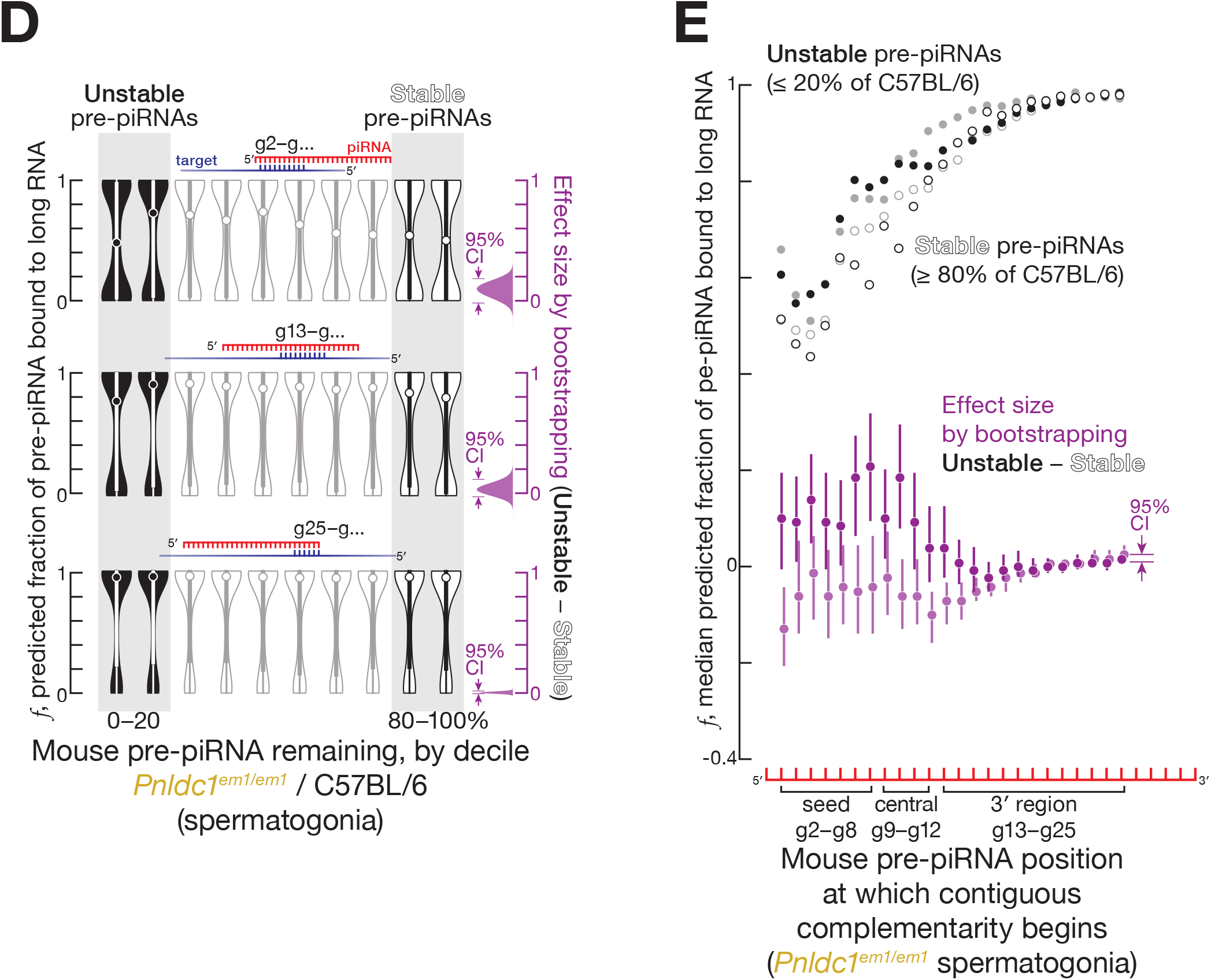
**Related to Figure 5.** (A) Median (*n* = 3) abundance of piRNAs and pre-piRNAs from FACS-purified mouse primary spermatocytes, secondary spermatocytes, and round spermatids. RNA sizes used for analysis were: 24–33-nt small RNAs for C57BL/6 and *Henmt1*^*em1/em1*^, and 24– 45-nt small RNAs for *Pnldc1*^*em1/em1*^. (B) Predicted fraction of mouse pre-piRNAs bound via different small RNA regions to complementary sites in the transcriptome. A representative experiment is shown for mouse pachytene pre-piRNAs from FACS-purified *Pnldc1*^*em1/em1*^ primary spermatocytes. The 95% confidence interval for the effect size of median difference was calculated with 10,000 bootstrapping iterations. (C) Analysis of mouse pre-piRNAs from FACS-purified primary spermatocytes showing median predicted fraction bound for complementarity starting at pre-piRNA positions g2–g25 for stable (≥ 80% of C57BL/6 levels in *Pnldc1*^*em1/em1*^) and unstable pre-piRNAs (≤ 20% of C57BL/6 levels in *Pnldc1*^*em1/em1*^), as well as the difference between the two (i.e., unstable pre-piRNAs − stable pre-piRNAs). Two independent experiments are shown in different shades of the same color. The 95% confidence interval for the effect size of median difference was calculated with 10,000 bootstrapping iterations. (D) Predicted fraction of mouse pre-piRNAs bound via different small RNA regions to complementary sites in the transcriptome. Data are from a representative experiment for mouse pre-pachytene pre-piRNAs from FACS-purified spermatogonia. The 95% confidence interval for the effect size of median difference was calculated with 10,000 bootstrapping iterations. (E) Analysis of mouse pre-piRNAs from FACS-purified spermatogonia showing median predicted fraction bound for complementarity starting at pre-piRNA positions g2–g25 for stable (≥ 80% of C57BL/6 levels in *Pnldc1*^*em1/em1*^) and unstable pre-piRNAs (≤ 20% of C57BL/6 levels in *Pnldc1*^*em1/em1*^), as well as the difference between the two (i.e., unstable pre-piRNAs − stable pre-piRNAs). Two independent experiments are shown in different shades of the same color. The 95% confidence interval for the effect size of median difference was calculated with 10,000 bootstrapping iterations.

**Figure S6.**
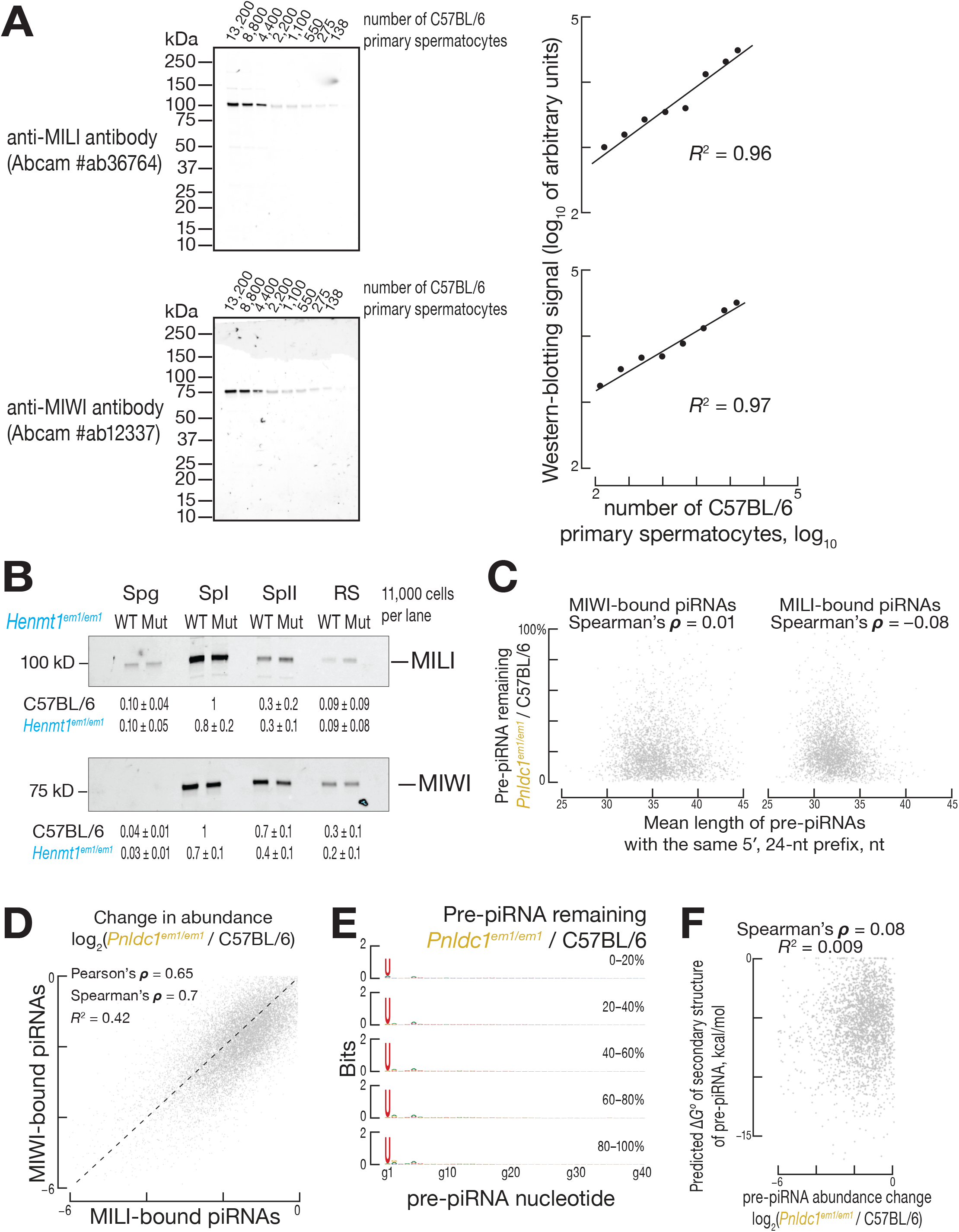
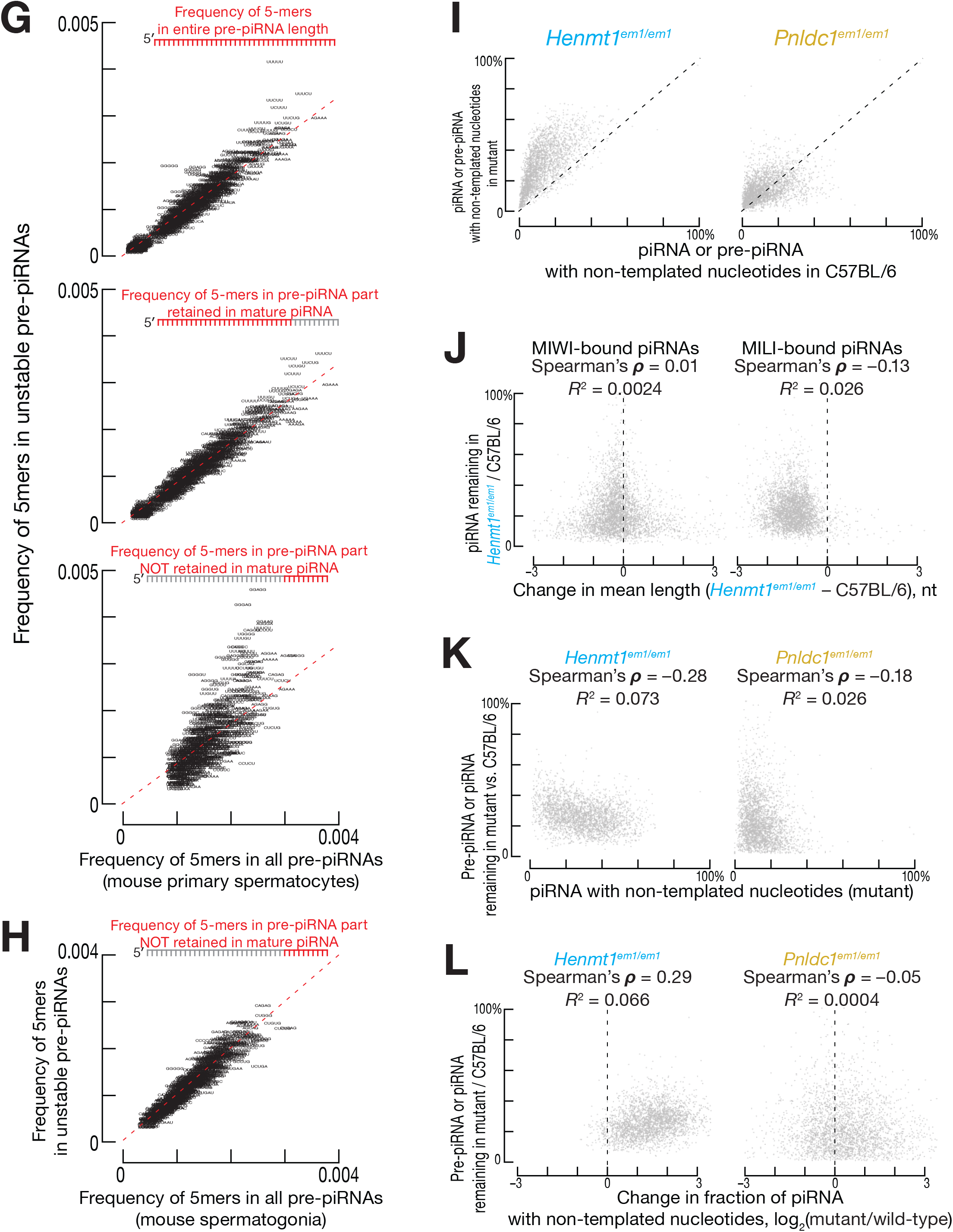
**Related to Figure 5.** (A) Dynamic range of western-blotting assay using anti-MILI (Abcam #ab36764) and anti-MIWI (Abcam Cat# ab12337) antibodies. (B) Relative abundance of MILI and MIWI (mean ± SD, *n* = 3) in C57BL/6 and *Henmt1*^*em1/em1*^ animals. FACS-purified cell types: Spg, spermatogonia; SpI, primary spermatocytes; SpII, secondary spermatocytes; RS, round spermatids. Each lane contains lysate from ~11,000 cells. (C) Length and stability of pre-piRNAs from FACS-purified *Pnldc1*^*em1/em1*^ primary spermatocytes. (D) Change in abundance for MILI- and MIWI-bound pre-piRNAs in FACS-purified primary spermatocytes from *Pnldc1*^*em1/em1*^ mice. (E) Positional nucleotide bias of pre-piRNAs in FACS-purified *Pnldc1*^*em1/em1*^ primary spermatocytes. (F) Predicted ΔG° of secondary structure and the change in abundance for pre-piRNAs in FACS-purified primary spermatocytes of *Pnldc1*^*em1/em1*^ mice. (G) Frequency of 5-mers in pre-piRNA sequences in FACS-purified *Pnldc1*^*em1/em1*^ mouse primary spermatocytes. (H) Frequency of 5-mers in pre-piRNA sequences in FACS-purified *Pnldc1*^*em1/em1*^ mouse spermatogonia. (I) Change in the fraction of piRNAs and pre-piRNAs bearing 3′ terminal non-templated nucleotides in FACS-purified *Henmt1*^*em1/em1*^ and *Pnldc1*^*em1/em1*^ primary spermatocytes. (J) Stability and change in mean length for piRNAs from FACS-purified *Henmt1*^*em1/em1*^ primary spermatocytes. (K) Stability and fraction bearing 3′ terminal non-templated nucleotides for piRNAs from *Henmt1*^*em1/em1*^ and pre-piRNAs from *Pnldc1*^*em1/em1*^ FACS-purified primary spermatocytes. (L) Stability and change in fraction bearing 3′ terminal non-templated nucleotides for piRNAs in *Henmt1*^*em1/em1*^ and pre-piRNAs in *Pnldc1*^*em1/em1*^ FACS-purified primary spermatocytes.

**Figure S7.**
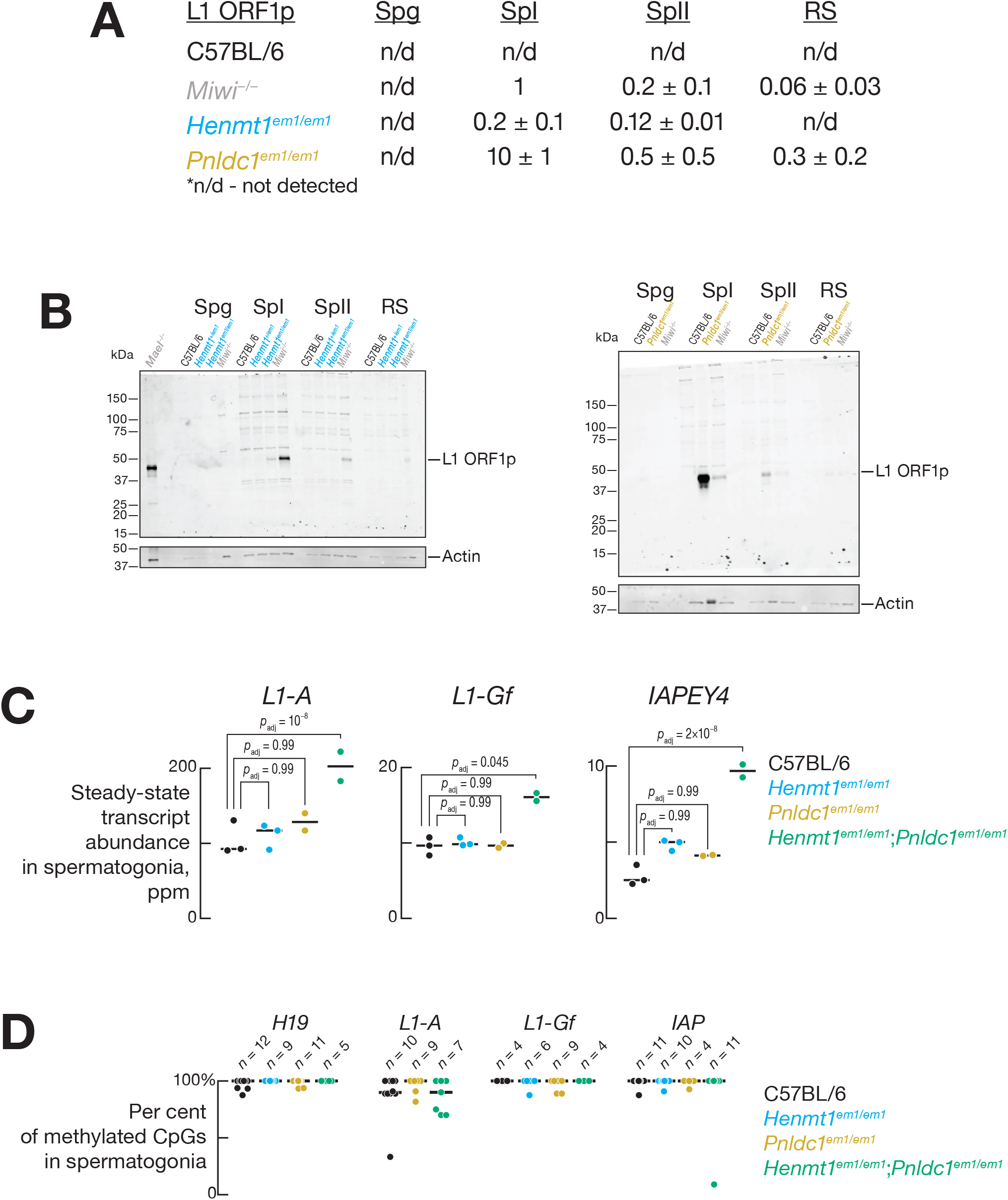
**Related to Figure 6.** (A) Relative abundance of L1 ORF1 protein (L1 ORF1p) in C57BL/6, *Miwi^—/—^*, *Henmt1*^*em1/em1*^ and *Pnldc1*^*em1/em1*^ animals (mean ± SD; *n* = 2 for *Pnldc1*^*em1/em1*^; *n* = 3 for *Henmt1*^*em1/em1*^). FACS-purified cell types: Spg, spermatogonia; SpI, primary spermatocytes; SpII, secondary spermatocytes; RS, round spermatids. Data were normalized to *Miwi*^−/−^ primary spermatocytes. Representative western blotting images used for quantification are shown in (B). (B) Relative abundance of L1 ORF1p in FACS-purified male germ cells of C57BL/6, *Henmt1*^*em1/em1*^ and *Pnldc1*^*em1/em1*^ mice assessed by Western blotting. Spg, spermatogonia; SpI, primary spermatocytes; SpII, secondary spermatocytes; RS, round spermatids. Each lane contains lysate from ~27,000 cells. *Miwi*^−/−^ and *Mael^−/−^* cells provided positive controls. (C) Relative steady-state abundance of transposon transcripts in spermatogonia from *Henmt1*^*em1/em1*^, *Pnldc1*^*em1/em1*^, and *Henmt1*^*em1/em1*^; *Pnldc1*^*em1/em1*^ mice. Adjusted *p* values were corrected for multiple hypothesis testing using the Benjamini-Hochberg procedure. (D) CpG methylation levels of transposons in spermatogonia of *Henmt1*^*em1/em1*^, *Pnldc1*^*em1/em1*^, and *Henmt1*^*em1/em1*^; *Pnldc1*^*em1/em1*^ mice. The number of individual clones is shown. (E) Predicted fraction of mouse pre-piRNAs bound via different small RNA regions to complementary sites in the transcriptome. Data are from a representative experiment for mouse pre-pachytene pre-piRNAs from FACS-purified *Henmt1*^*em1/em1*^; *Pnldc1*^*em1/em1*^ spermatogonia. The 95% confidence interval for the effect size of median difference was calculated with 10,000 bootstrapping iterations. (F) Analysis of mouse pre-piRNAs from FACS-purified spermatogonia showing median predicted fraction bound for complementarity starting at pre-piRNA positions g2–g25 for stable (≥ 80% of C57BL/6 levels in *Henmt1*^*em1/em1*^; *Pnldc1*^*em1/em1*^) and unstable pre-piRNAs (≤ 20% of C57BL/6 levels in *Henmt1*^*em1/em1*^; *Pnldc1*^*em1/em1*^), as well as the difference between the two (i.e., unstable pre-piRNAs − stable pre-piRNAs). Two independent experiments are shown in different shades of the same color. The 95% confidence interval for the effect size of median difference was calculated with 10,000 bootstrapping iterations.

## SUPPLEMENTAL ITEM TITLES

**Table S1. Transposon Sub-families Derepressed in Spermatogonia of***Henmt1*^*em1/em1*^;*Pnldc1*^*em1/em1*^ **Mice and in Primary Spermatocytes from***Henmt1*^*em1/em1*^ **and***Pnldc1*^*em1/em1*^ **Mice, Related to Figure 5**.

**Table S2. mRNAs whose Derepression in***Henmt1*^*em1/em1*^ **and***Pnldc1*^*em1/em1*^ **Mice can be Explained by the Reduced Cleavage by Destabilized piRNAs, Related to Figure 5.**

Data are for the intersection of datasets from two biological samples.

**Table S3. Sequences of Synthetic 5′ Monophosphorylated Spike-in RNA Oligonucleotides Included in the Mix for Small RNA Sequencing Libraries.**

**Table S4. Number of Cells and Amount of Spike-In Mix Used to Prepare Small RNA Sequencing Libraries, Related to Figure 1.**

**Table S5. Number of Cells and Amount of ERCC Spike-In Mix 1 Used to Prepare RNA Sequencing Libraries (A) and List of Libraries of 5′ Monophosphorylated Long RNAs (B), Related to Figure 1.**

**Table S6. Primers Used for Bisulfite Analysis of DNA Methylation, Related to Figure S7.**

**Table S7. Data Sets Used in This Study, Related to Figures 2, 3, and 4.**

## STAR METHODS

### RESOURCE AVAILABILITY

#### Lead Contact

Further information and requests for resources and reagents should be directed to, and will be fulfilled by, the Lead Contact, Phillip D. Zamore, or by completing the request form at https://www.zamorelab.umassmed.edu/reagents.

#### Materials Availability

Strains generated in this study are available for non-commercial use upon request without restriction by request or, where indicated, from the Bloomington Drosophila Stock Center (https://bdsc.indiana.edu) or the Jackson Laboratory (https://www.jax.org/jax-mice-and-services/find-and-order-jax-mice).

#### Data and Code Availability

Sequencing data are available from the National Center for Biotechnology Information Sequence Read Archive using accession number PRJNA660633.

## EXPERIMENTAL MODEL AND SUBJECT DETAILS

### Mouse Strains and Mutants

Mice (wild-type C57BL/6J, IMSR Cat# JAX:000664, RRID:IMSR_JAX:000664; and *Pnldc1*^*em1Pdz/em1Pdz*^ mutants, MGI: 6161374) were maintained and sacrificed according to the guidelines of the Institutional Animal Care and Use Committee of the University of Massachusetts Medical School.

Guide RNA (sgRNA: 5′-GGC ATC TCC ACA TCC CAG GTC GG-3′) targeting exon 4 of *Henmt1* to generate *Henmt1*^*em1Pdz/em1Pdz*^ (MGI: 6452642) was designed using CRISPR design tool (crispr.mit.edu/). sgRNA was transcribed with T7 RNA Polymerase and then purified by electrophoresis on 10% denaturing polyacrylamide gel. As a donor to generate *Henmt1*^*em1Pdz/em1Pdz*^ single stranded oligonucleotide was ordered from IDT. A mix of sgRNA (20 ng/μl), Cas9 mRNA (50 ng/μl, TriLink Biotechnologies, L-7206) and 195-nt, single-stranded oligonucleotide (100 ng/μl) donor were injected together into the pronucleus of one-cell C57BL/6 zygotes in M2 medium (Sigma, M7167). After injection, the zygotes were cultured in KSOM with amino acids at 37°C under 5% CO2 until the blastocyst stage (3.5 days), which further transferred into uterus of pseudopregnant ICR females at 2.5 dpc.

To screen for mutant founders, genomic DNA extracted from tail tissues was analyzed by PCR. Primers used for genotyping *Henmt1*^*em1Pdz/em1Pdz*^ are 5′-GTT GCC AAC GCT GTA GCC-3′ and 5′-AAT AAG GGC ACC CTG CAC TA-3′. Mutant sequences were confirmed by Sanger sequencing. In *Henmt1*^*em1Pdz/em1Pdz*^ mutants, genomic sequence is altered resulting in changing amino acid residues 54–58 from DLGCG to NAVAV, which likely leads to loss of catalytic activity {Kirino and Mourelatos, 2007, #68317} and misfolding of the protein: *Henmt1* mRNA level in *Henmt1*^*em1Pdz/em1Pdz*^ is identical to that in C57BL/6 animals; HENMT1 protein level in *Henmt1*^*em1Pdz/em1Pdz*^ is ~0.04 of that in C57BL/6 animals (data not shown).

### *Drosophila melanogaster* Strains and Mutants

Fly stocks were maintained at 25°C. All strains were in the *w*^*1118*^ background. Before dissection, flies were isolated 0–3 days after eclosion and given yeast paste for two days. Fly ovaries or heads were then dissected and collected in 1× phosphate-buffered saline [pH 7.4] (1×PBS: 137 mM NaCl, 2.7 mM KCl, 10 mM Na_2_HPO_4_, 1.8 mM KH_2_PO_4_) cooled on ice. Ovaries or heads were washed once with ice-cold 1×PBS and then used for subsequent experiments.

## METHOD DETAILS

### Mouse Phenotypic Analysis

All potential mutant male mice (2–8 months old) were housed with one 2–4 months old C57BL/6J female. The presence of a vaginal plug was examined the following morning to confirm insemination. If a plug was observed, the female was separated and observed for potential pregnancy. Males mated to females who failed to produce pups within 2 months after a vaginal plug was detected were deemed sterile. Presence of epididymal sperm, testis weight, and testis histology were also scored.

#### Histology

Testes were (1) collected from 2–6 month-old mice; (2) fixed overnight in Bouin’s solution; (3) washed three times with 70% (v/v) ethanol then stored in 70% ethanol. Tissues were embedded in paraffin and cut into 5 μm cross-sections, then stained with hematoxylin and counter stained with eosin (UMass Morphology Core).

### FACS Isolation and Immunostaining of Mouse Germ Cells

Testes of 2–5 month-old mice were isolated, decapsulated, and incubated for 15 min at 33°C in 1× Gey′s Balanced Salt Solution (GBSS, Sigma, G9779) containing 0.4 mg/ml collagenase type 4 (Worthington LS004188) rotating at 150 rpm. Seminiferous tubules were then washed twice with 1× GBSS and incubated for 15 min at 33°C in 1× GBSS with 0.5 mg/ml Trypsin and 1 μg/ml DNase I rotating at 150 rpm. Next, tubules were homogenized by pipetting through a glass Pasteur pipette for 3 min on ice. Fetal bovine serum (FBS; 7.5% f.c., v/v) was added to inactivate trypsin, and the cell suspension was then strained through a pre-wetted 70 μm cell strainer and cells collected by centrifugation at 300 × *g* for 10 min. The supernatant was removed, cells were resuspended in 1× GBSS containing 5% (v/v) FBS, 1 μg/ml DNase I, and 5 μg/ml Hoechst 33342 (Thermo Fisher, 62249) and rotated at 150 rpm for 45 min at 33°C. Propidium iodide (0.2 μg/ml, f.c.; Thermo Fisher, P3566) was added, and cells strained through a single pre-wetted 40 μm cell strainer. Four-way cell sorting (spermatogonia, primary spermatocytes, secondary spermatocytes, round spermatids; File S1) using a FACSAria II Cell Sorter (BD Biosciences; UMass Medical School FACS Core) was performed as described {Bastos et al., 2005, #41953} with modifications. Briefly, the 355-nm laser was used to excite Hoechst 33342; the 488-nm laser was used to excite Propidium iodide and record forward (FSC) and side (SSC) scatter. Propidium iodide emission was detected using a 610/20 bandpass filter (YG PE-Texas Red-A in File S1). Hoechst 33342 emission was recorded using 450/50 (UV-B Blue DAPI-A in File S1) and 670/50 (UV-A Red Side Pop-A in File S1) band pass filters.

Germ cell stages in the unsorted population and the purity of sorted fractions were assessed by immunostaining aliquots of cells. Cells were incubated for 20 min in 25 mM sucrose and then fixed on a slide with 1% (w/v) paraformaldehyde containing 0.15% (v/v) Triton X-100 for 2 h at room temperature in a humidifying chamber. Slides were washed sequentially for 10 min in: (1) 1× PBS containing 0.4% (v/v) Photo-Flo 200 (Kodak, 1464510); (2) 1× PBS containing 0.1% (v/v) Triton X-100; and (3) 1× PBS containing 0.3% (w/v) BSA, 1% (v/v) donkey serum (Sigma, D9663), and 0.05% (v/v) Triton X-100. After washing, slides were incubated with primary antibodies in 1× PBS containing 3% (w/v) BSA, 10% (v/v) donkey serum, and 0.5% (v/v) Triton X-100 overnight at room temperature in a humidified chamber. Rabbit polyclonal anti-SYCP3 (Abcam Cat# ab15093, RRID:AB_301639, 1:1000 dilution) and mouse monoclonal anti-γH2AX (Millipore Cat# 05-636, RRID:AB_309864, 1:1000 dilution) were used as primary antibodies. Slides were washed again as described and then incubated with secondary donkey anti-mouse IgG (H+L) Alexa Fluor 594 (Thermo Fisher Scientific Cat# A-21203, RRID:AB_2535789, 1:2000 dilution) or donkey anti-rabbit IgG (H+L) Alexa Fluor 488 (Thermo Fisher Scientific Cat# A-21206, RRID:AB_2535792, 1:2000 dilution) antibodies for 1 h in at room temperature in a humidified chamber. After incubation, slides were washed three times (10 min each) in 1× PBS containing 0.4% (v/v) Photo-Flo 200 and once for 10 min in 0.4% (v/v) Photo-Flo 200. Finally, slides were dried, mounted with ProLong Gold Antifade Mountant with DAPI (Thermo Fisher, P36931). To assess the purity of sorted fractions, 50–100 cells were staged by DNA, γH2AX, and SYCP3 staining {Bastos et al., 2005, #41953}. All samples met these criteria:

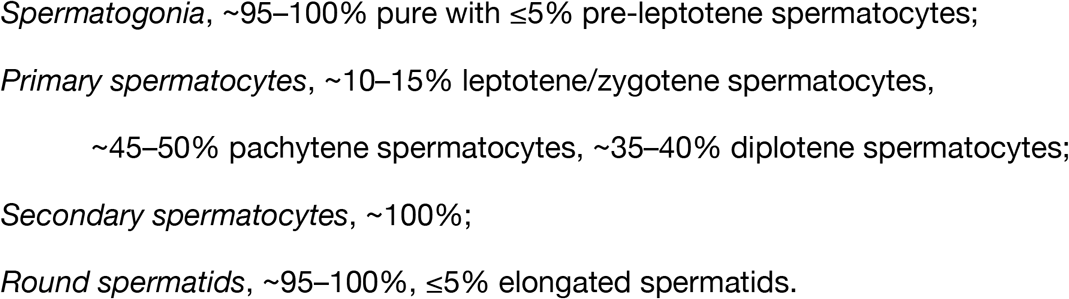

### Western Blotting

Cells were homogenized in Lysis Buffer (20 mM Tris-HCl pH 7.5, 2.5 mM MgCl2, 200 mM NaCl, 0.05% (v/v) NP-40, 0.1 mM EDTA, 1 mM 4-(2-Aminoethyl) benzenesulfonyl fluoride hydrochloride, 0.3 μM Aprotinin, 40 μM Bestatin, 10 μM E-64, 10 μM Leupeptin) and centrifuged at 20,000 × *g* for 20 min at 4°C. The supernatant was moved to a new tube, an equal volume of loading dye (120 mM Tris-HCl, pH 6.8, 4% (w/v) SDS, 20% (v/v) glycerol, 2.5% (v/v) 2-Mercaptoethanol, 0.2% (w/v) bromophenol blue) was added, and the sample incubated at 90°C for 5 min and resolved by electrophoresis through a 4–20% gradient polyacrylamide/SDS gel (Bio-Rad Laboratories, 5671085). Next, proteins were transferred to PVDF (Millipore, IPVH00010), the membrane blocked in Blocking Buffer (Rockland Immunochemicals, MB-070) at room temperature for 2 h and then incubated overnight at 4°C in Blocking Buffer containing primary antibody (anti-mouse PIWIL2/MILI, Abcam Cat# ab36764, RRID:AB_777284, 1:1000 dilution; anti-PIWIL1/MIWI, Abcam Cat# ab12337, RRID:AB_470241, 1:1000 dilution; anti-mouse LINE-1 ORF1p rabbit polyclonal, 1:10000 dilution (generous gift of Alex Bortvin; {*Martin, 1991, #45866; Soper et al., 2008, #89272}). The membrane was washed three times (30 min each) with Blocking Buffer at room temperature and incubated for 2 h at room temperature with donkey anti-rabbit IRDye 680RD secondary antibody (LI-COR Biosciences Cat# 926-68073, RRID:AB_1095444, diluted 1:20,000) in Blocking Buffer. Then the membrane was washed three times (30 min each) with Blocking Buffer at room temperature and the signal detected using an Odyssey Infrared Imaging System. In our assays, anti-MILI and anti-MIWI assesses the abundance of MILI and MIWI across a broad dynamic range: 1% to 120% of the level in C57BL/6 primary spermatocytes (Figure S6A).

### Small RNA Immunoprecipitation

Sorted mouse germ cells were homogenized with Lysis Buffer (see Western Blotting) and then centrifuged at 20,000 × *g* for 20 min at 4°C, retaining the supernatant. Anti-MIWI (Wako, Cat# 017-23451, RRID:AB_2721829, ~5 μg) or anti-MILI (Abcam Cat# ab36764, RRID:AB_777284, ~5 μg) antibodies were incubated with rotation with 30 μl of Protein G Dynabeads (Thermo Fisher, 10003D) in 1× PBS containing 0.02% (v/v) Tween 20 (PBST) at 4°C for 1 h. The bead-antibody complex was washed with PBST. Freshly prepared testis or cell lysate was added to the bead-antibody complex and incubated with rotation at 4°C overnight. The next day, the beads were washed once with lysis buffer and three times with 0.1 M trisodium citrate. After washing, RNA was extracted with Trizol reagent (Thermo Fisher, 15596026) and used for small RNA library preparation.

### Small RNA-seq Library Preparation

Total RNA from sorted mouse germ cells was extracted using the mirVana miRNA isolation kit (Thermo Fisher, AM1560). Small RNA libraries were constructed as described {Gainetdinov et al., 2018, #64592} with several modifications. Briefly, before library preparation, a mix of nine synthetic RNA oligonucleotides (Table S3) was added to each RNA sample to enable absolute quantification of small RNAs (Table S4; median cell volume from {*Gainetdinov et al., 2018, #64592} was used to calculate intracellular concentrations). To reduce ligation bias and eliminate PCR duplicates, 3′ and 5′ adaptors each contained nine random nucleotides at their 5′ and 3′ ends, respectively {Fu et al., 2018, #88843}; 3′ adaptor ligation reactions contained 20% (w/v) PEG-8000 (f.c.). After 3′ adaptor ligation, RNA was purified by 15% urea polyacrylamide gel electrophoresis (PAGE), selecting for 15–55 nt small RNAs (i.e., 50– 90 nt with 3′ adaptor). Small RNA-seq libraries for 2–4 biological samples were sequenced together using a NextSeq 500 (Illumina) to obtain 75 nt, single-end reads. Data sets of MILI- and MIWI-bound piRNAs in C57BL/6 and *Pnldc1*^*em1Pdz/em1Pdz*^ are from {*Gainetdinov et al., 2018, #64592}.

### RNA-seq Library Preparation

Total RNA from sorted germ cells was extracted using the mirVana miRNA isolation kit (Thermo Fisher, AM1560) and used for library preparation as described {Zhang et al., 2012, #98422} with modifications, including the addition of the ERCC spike-in mix to enable absolute quantification of RNAs and the use of unique molecular identifiers to eliminate PCR duplicates {Fu et al., 2018, #88843}. Briefly, before library preparation, 1 μl of 1:100 dilution of ERCC spike-in mix 1 (Thermo Fisher, 4456740, LOT00418382; Table S5) was added to 0.5–1 μg total RNA to enable absolute quantification of mRNA. For ribosomal RNA depletion, RNA was hybridized in 10 μl to a pool of 186 rRNA antisense oligos (0.05 μM each) in 10 mM Tris-HCl (pH 7.4), 20 mM NaCl, heating the mixture to 95°C, cooling it at −0.1°C/sec to 22°C, and incubating at 22°C for 5 min. RNase H (10U; Lucigen, H39500) was added and the mixture incubated at 45°C for 30 min in 20 μl containing 50 mM Tris-HCl (pH 7.4), 100 mM NaCl, and 20 mM MgCl2. Reaction was adjusted to 50 μl with 1× TURBO DNase buffer (ThermoFisher) and then incubated with 4U DNase (Thermo Fisher, AM2238) at 37°C for 20 min. Next, RNA was purified using RNA Clean & Concentrator-5 (Zymo Research, R1016). RNA-seq libraries for three samples were sequenced together using a NextSeq 500 (Illumina) to obtain 79 + 79 nt, paired-end reads. Data sets of C57BL/6 secondary spermatocytes and round spermatids are from {*Gainetdinov et al., 2018, #64592}.

### Cloning and Sequencing of 5′ Monophosphorylated Long RNAs

Total RNA from sorted mouse germ cells was extracted using mirVana miRNA isolation kit (Thermo Fisher, AM1560) and used to prepare a library of 5′ monophosphorylated long RNAs as described {Wang et al., 2014, #4083}. Libraries for two independent biological replicates were sequenced using a NextSeq 500 (Illumina) to obtain 79 + 79 nt, paired-end reads (Table S5).

### DNA Methylation Detection

DNA methylation was assessed using DNA bisulfate sequencing. FACS-sorted spermatogonia were lysed and DNA was treated with bisulfite using EZ-DNA Methylation Direct Kit (Zymo Research). Imprinted locus H19 was used as the methylated DNA control. Bisulfate-treated DNA served as the template in one round (L1-Gf and L1-A) or two nested rounds (H19 and IAP) of PCR with EipMark Hot Start *Taq* DNA Polymerase (NEB) using the following protocol: initial denaturation – 95°C for 30 seconds; 35 cycles of 95°C for 30 seconds, annealing temperature for 60 seconds, and 68°C for 30 seconds; final extension – 68°C for 5 minutes (Table S6 contains primer sequences and annealing temperatures). Primers were specific to different genomic copies of the same transposon family. PCR products were cloned and sequenced.

## QUANTIFICATION AND STATISTICAL ANALYSIS

### Analysis of Small RNA Data Sets

The sequences were filtered by requiring their Phred quality score to be ≥ 20 for all nucleotides, the 3′ adapter and PCR duplicates were removed from raw reads. Sequences of synthetic spike-in oligonucleotides were identified allowing no mismatches (Table S3). Reads not fully matching the genome were analyzed using the Tailor pipeline {Chou et al., 2015, #600} to account for non-templated tailing of small RNAs. All unambiguously mapping piRNA or pre-piRNA reads were grouped by their 5′, 24-nt prefix.

### RNA-seq Library Analysis

RNA-seq analysis was performed using piPipes for genomic alignments {Han et al., 2015, #98844} and custom scripts to remove PCR duplicates {Fu et al., 2018, #88843}. Briefly, sequences were first reformatted to extract unique molecular identifiers {Fu et al., 2018, #88843}, and then aligned to ribosomal RNA using Bowtie2 (v2.2.0; {*Langmead and Salzberg, 2012, #26610}). Unaligned reads were then mapped to mouse genome mm10 using STAR (v2.3.1; {*Dobin et al., 2013, #39763}), and PCR duplicates removed {Fu et al., 2018, #88843}. Sequencing depth and gene quantification were calculated with StringTie (v1.3.4; {*Pertea et al., 2016, #52019}). Differential expression analysis was performed using DESeq2 (v1.18.1; {*Love et al., 2014, #2502}). In parallel, reformatted reads were aligned to an index of ERCC spike-in transcripts (Thermo Fisher, 4456740, LOT00418382) using Bowtie (v1.0.0; {*Langmead et al., 2009, #34306}), PCR duplicates were removed, and the absolute quantity of transcripts assessed (Table S5).

### Analysis of 5′ Monophosphorylated Long RNA Sequencing Data

Analysis of 5′ monophosphorylated long RNA sequencing data was performed with piPipes {Han et al., 2015, #98844}. Briefly, RNAs were first aligned to ribosomal RNA (rRNA) sequences using Bowtie2 (v2.2.0). Unaligned reads were then mapped to mouse genome mm10 using STAR (v2.3.1) and alignments with soft clipping of ends were removed with SAMtools (v1.0.0; {*Li et al., 2009, #15771}).

### Biochemical Model to Calculate Fraction of Small RNA Bound to Long RNA

We considered the enzymatic mechanism:

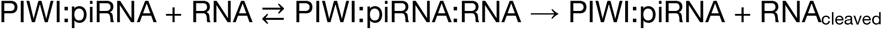

For a given piRNA binding site *i*, we considered these molecular species,

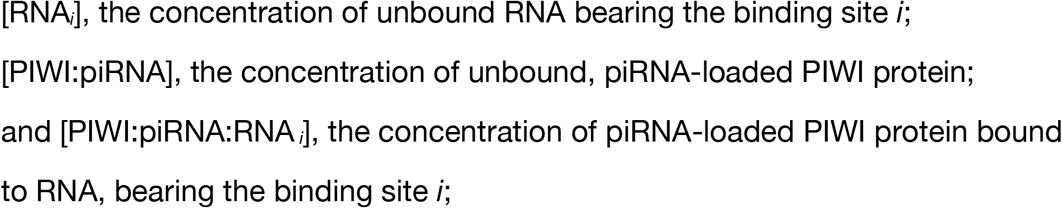

and defined the following kinetic rates,

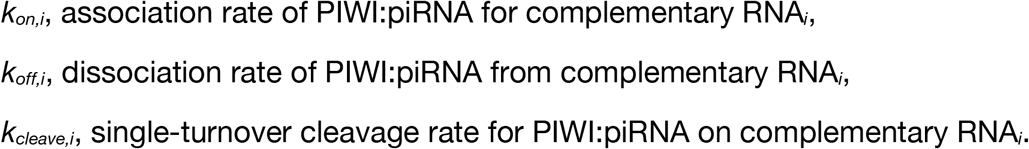

This rate equation describes PIWI:piRNA activity:

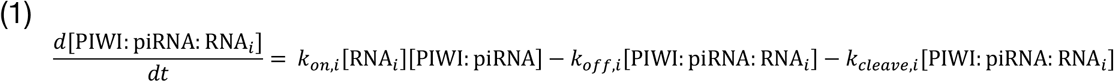

Assuming that the cleavage step is slow, i.e., *kcleave* ≪ *koff*, equation (1) becomes:

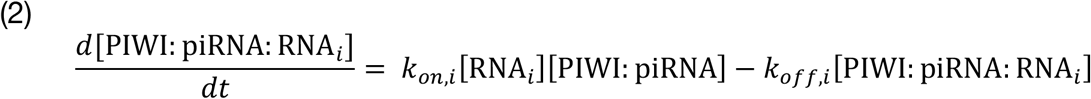

At equilibrium (i.e., when *d*[PIWI:piRNA:RNA*i*]/*dt* is 0), we can rewrite equation (2) as:

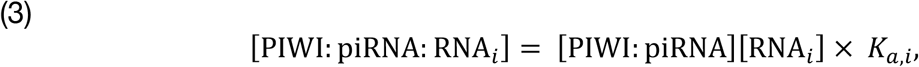

where *K_a_* represents the association constant of PIWI:piRNA for complementary RNA.

We define fraction bound *f* as:

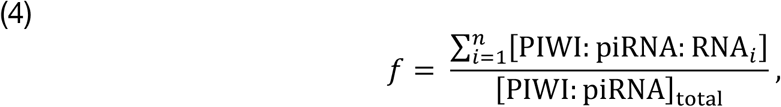

where *n* is the total number of piRNA binding sites.

Substituting [PIWI:piRNA:RNA*i*] in equation (4) using equation (3) yields

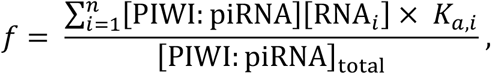

which rearranges to

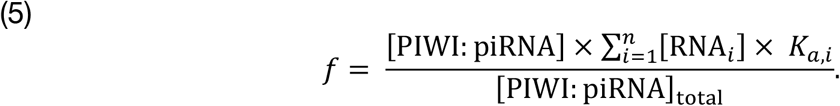

Considering that

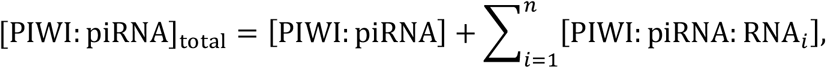

equation (5) becomes:

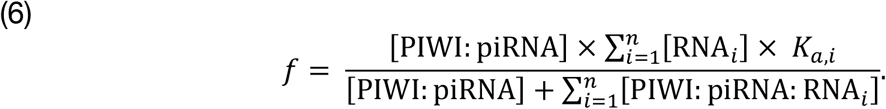

Substituting [PIWI:piRNA:RNA *i*] in equation (6) using equation (3) yields

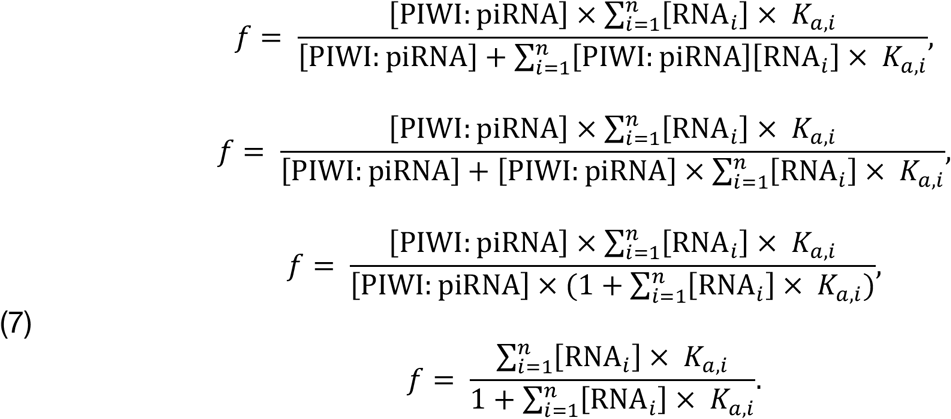

Using the assumptions (1) that the rank order of [RNA] can be approximated by the rank order of [RNA]_total_, and (2) that the rank order of affinities of PIWI:piRNA for complementary RNA can be approximated using the computationally predicted Gibbs

free energy (Δ*G*°) of base pairing between two RNA strands 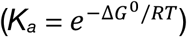, equation (7) becomes equation (8):

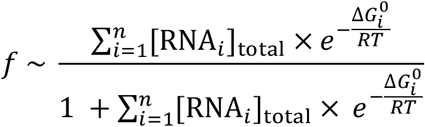

where *R* = 1.987 cal∙K^−1^∙mol^−1^ and *T* = 298.15 K for fly heads, *T* = 300.15 K for S2 cells, *T* = 306.15 K for mouse testis, and *T* = 310.15 K for mESC and MEF cells.

PredictedΔ*G*° was calculated from nearest neighbor values using RNAfold 2.4.14 {Lorenz et al., 2011, #46902}. Total intracellular concentrations of long RNAs ([RNA*i*]total) in mouse germ cells were measured with RNA-seq using ERCC spike-ins (Table S5) and cellular volumes reported from {*Gainetdinov et al., 2018, #64592}. For other types of data (Table S7), relative transcript abundance was converted to intracellular concentration based on the mean total transcript concentration in mouse spermatogonia and primary spermatocytes (~1,500,000 transcripts per 1000 μm^3^). For fly piRNAs, the mean (*n* = 2) change of piRNA abundance and the mean (*n* = 2) abundance of mRNAs were used to calculate a fraction bound estimate for each region of each piRNA. For fly and mouse miRNAs, the mean (*n* = 2) miRNA half-life and the mean (*n* = 2) abundance of mRNAs were used to calculate a fraction bound estimate for each region of each miRNA.

### Statistical Tests

Statistical tests are described in the figure legends.

