## Supplementary material for "Terminal Modification, Sequence, and Length Determine Small RNA Stability in Animals": Tables S1, S3, S6, and S7

**Table S1. Transposon Sub-families Derepressed in Spermatogonia of *Henmt1*$^{em1/em1}$; *Pnldc1*$^{em1/em1}$ Mice and in Primary Spermatocytes from *Henmt1*$^{em1/em1}$ and *Pnldc1*$^{em1/em1}$ Mice, Related to Figure S7.**

(A) Primary Spermatocytes from *Henmt1*$^{em1/em1}$ mice.

| Transposon subfamily | Change in steady-state transcript abundance in primary spermatocytes, $\log_2$ (*Henmt1*$^{em1/em1}$ / C57BL/6 | $p_{adj}$ |
|---|---|---|
| MER67D | -1.14 | 0.05 |
| MER90a | -0.90 | 0.04 |
| MamGyp-int | -0.70 | 0.10 |
| MER45A | -0.68 | 0.07 |
| MurERV4-int | 0.70 | 0.08 |
| RLTR44-int | 0.80 | 0.03 |

(B) Primary Spermatocytes from *Pnldc1*$^{em1/em1}$ mice.

| Transposon subfamily | Change in steady-state transcript abundance in primary spermatocytes, $\log_2$ (*Pnldc1*$^{em1/em1}$ / C57BL/6) | $p_{adj}$ |
|---|---|---|
| MER90a | -1.22 | 0.05 |
| LTR28B | -0.97 | 0.05 |
| X7A_LINE | -0.92 | 0.05 |
| RMER17A-int | -0.90 | 0.09 |
| MER67C | -0.80 | 0.05 |
| L1MC | -0.50 | 0.10 |
| MLT1E1 | 0.54 | 0.10 |
| RLTR14 | 0.56 | 0.07 |
| RLTR44-int | 0.56 | 0.06 |
| L1Md_T | 0.58 | 0.01 |
| RMER15-int | 0.60 | 0.05 |
| L1MEh | 0.68 | 0.09 |
| LTR67B | 0.70 | 0.00 |
| IAPLTR3 | 0.79 | 0.03 |
| RLTR10B | 0.89 | 0.00 |
| IAPLTR3-int | 0.90 | 0.00 |
| RLTR20A2B_MM | 0.91 | 0.06 |
| Charlie18a | 0.96 | 0.05 |
| RLTR24 | 0.96 | 0.00 |
| LTR102_Mam | 0.97 | 0.00 |
| MLT1E | 0.97 | 0.00 |
| L1Md_A | 1.23 | 0.00 |

(C) Spermatogonia from *Henmt1*$^{em1/em1}$; *Pnldc1*$^{em1/em1}$ mice.

| Transposon subfamily | Steady-state transcript abundance change in spermatogonia; $\log_2$ of *Henmt1*$^{em1/em1}$; *Pnldc1*$^{em1/em1}$/ C57BL/6 | $p_{adj}$-value |
|---|---|---|
| L1Md_Gf | 0.79 | 0.045 |
| L1Md_A | 1.00 | 0.00000001 |
| RLTR4_MM-int | 1.20 | 0.0003 |
| IAPEy-int | 1.40 | 0.0054 |
| IAPEY4_I-int | 1.86 | 0.00000002 |
| IAPEY5_I-int | 2.28 | 0.074 |

**Table S3. Sequences of Synthetic 5′ Monophosphorylated Spike-in RNA Oligonucleotides Included in the Mix for Small RNA Sequencing Libraries.**

| Sequence |
|---|
| 5′-UGCUAGUCUGUUAUCGACCUGACCUCAUAG-3′ |
| 5′-UGCUAGUCUGUUCGAUACCUGACCUCAUAG-3′ |
| 5′-UGCUAGUCUGUUGUCACGAAGACCUCAUAG-3′ |
| 5′-UGCUAGUCUUAUCGACCUCCUCAUAG-3′ |
| 5′-UGCUAGUCUUCGAUACCUCCUCAUAG-3′ |
| 5′-UGCUAGUCUUGUCACGAACCUCAUAG-3′ |
| 5′-UGCUAGUUAUCGACCUUCAUAG-3′ |
| 5′-UGCUAGUUCGAUACCUUCAUAG-3′ |
| 5′-UGCUAGUUGUCACGAAUCAUAG-3′ |

**Table S6. Primers Used for Bisulfite Analysis of DNA Methylation, Related to Figure S7.**

| Name | Sequence | PCR product length, bp | Annealing temperature for cycles 1–10 (touchdown), ºC | Annealing temperature for cycles 11–35, ºC |
|---|---|---|---|---|
| H19_PCR1_Forward | GAGTATTTAGGAGGTATAAGAATT | 423 | 65–56 | 55 |
| H19_PCR1_Reverse | ATCAAAAACTAACATAAACCCCT | | | |
| H19_PCR2_Forward | GTAAGGAGATTATGTTTATTTTTGG | | | |
| H19_PCR2_Reverse | CCTCATTAATCCCATAACTAT | | | |
| L1-Gf_Forward | GTTAGAGAATTTGATAGTTTTTGGAATAGG | 270 | 65–56 | 55 |
| L1-Gf_Reverse | CCAAAACAAAACCTTTCTCAAACACTATAT | | | |
| L1-A_Forward | GGATTTTAAGATTTTTGGTGAGTGGAATATAG | 292 | 65–56 | 55 |
| L1-A_Reverse | ACTCTCTTCTTACAAAACAAATACCCAAATATCT | | | |
| IAP1d1_PCR1_Forward | GTTTGTAATGGTGGGAGAT | 259 | 60–51 | 50 |
| IAP1d1_PCR1_Reverse | ATTCTAAAATAAAATATCCCTCC | | | |
| IAP1d1_PCR2_Forward | AAATAAATTGTGGGAAGT | | | |
| IAP1d1_PCR2_Reverse | CAAAAAAAACACCACAAACCAAAAT | | | |

**Table S7. Data Sets Used in This Study, Related to Figures 2, 3, and 4.**

| Species | Accession Numbers | Small RNA Data Set Reference | Description |
|---|---|---|---|
| *Mus musculus* | SRR8862219, SRR8862220 | Genome Res. 2019; 29(11): 1777-1790 | Contact-Inhibited Mouse Embryonic Fibroblasts (RNA-seq) |
| *Mus musculus* | SRR8862221, SRR8862222 | Genome Res. 2019; 29(11): 1777-1790 | Dividing Mouse Embryonic Fibroblasts (RNA-seq) |
| *Mus musculus* | SRR8862223, SRR8862224 | Genome Res. 2019; 29(11): 1777-1790 | Mouse Embryonic Stem Cells (RNA-seq) |
| *Drosophila melanogaster* | SRR8877875 | Mol Cell. 2019; 75(4):756-768 | ago2 KO S2 Cells (RNA-seq) |
| *Drosophila melanogaster* | SRR1505815, SRR1505816 | Cell Rep. 2014; 9(6):2290-303 | GMR-gal4 x UAS-GFP ; GFP-positive cells (RNA-seq) |
